# Analgesic actions of Intrathecal NaV 1.7 antisense in rats: loss of antagonist channel binding, message depletion, and neuraxial distribution of oligonucleotide

**DOI:** 10.1101/2025.05.19.654680

**Authors:** Kaue Franco Malange, Julia Borges Paes Lemes, Briana Noble, Yuvicza Anchondo, Carlos Morado Urbina, Saee Jadhav, Sara Dochnal, Chiraag Kambalimath, Pedro Alvarez, Bethany Fitzsimmons, Curt Mazur, Holly B. Kordasiewicz, Kim Dore, Hien T. Zhao, Tony Yaksh

**Affiliations:** Department of Anesthesiology, School of Medicine, University of California San Diego, San Diego, USA; Department of Neuroscience, School of Medicine, University of California San Diego, San Diego, USA; IONIS Pharmaceuticals, Inc., Carlsbad, USA

**Keywords:** Nav 1.7, Channel binding, Dorsal Root Ganglia, Antisense, ATTO488PTx-II

## Abstract

**Background:** Genome targeting strategies to address NaV 1.7 mediated signaling in nociceptive afferents produce highly selective and persistent analgesic outcomes. Here, we analyze the concentration-dependent effects of the reduction of primary afferent NaV 1.7 channel expression by intrathecal delivery of an antisense oligonucleotide on pain behaviors and the covariance of *Scn9a* knock-down on NaV 1.7 message expression and channel binding.

**Methods:** Male Sprague-Dawley rats were implanted with lumbar intrathecal catheters and dosed with different NaV 1.7 ASO concentrations (100 to 3000 μg;10 μL). Pain behavior assays were conducted 0-28 days after ASO injections. Brain, spinal cords (SC) and dorsal root ganglia (DRGs) were collected. Quantification of ASO knock-down was assessed through RT-qPCR. NaV 1.7 expression was assessed by binding of NaV 1.7 fluorescent labeled antagonist (ATTO488PTx-II). Distribution studies were performed using anti-ASO antibody staining in brain, SCs and DRGs.

**Results:** NaV 1.7 message was detected in nerve, DRG and SC. Intrathecal ASO induced a concentration dependent gradient of knock down in DRGs (lumbar to cervical) of *Scn9a* mRNA and ATTO488PTx-II binding in small DRG neurons, and in spinal parenchyma, and a suppression of pain behaviors initiated by mechanical compression, inflammation and following intraplantar NaV1.7 agonist (OD1) or formalin. At 1000 µg, there was a 47% reduction in phase 2 flinching, a 60% reduction in DRG mRNA and a 36% reduction in ATTO488PTx-II DRG binding in comparison with mismatch controls. Although marked changes were seen at the sensory ganglia level and spinal dorsal horn, no changes in NaV 1.7 binding or mRNA were detected in sciatic nerves. Reduction in DRG message displayed a rostrocaudal gradient that corresponded with ASO distribution.

**Conclusions:** The study presents how NaV 1.7 ASOs reduce primary afferent channel binding through an effective knock-down on *Scn9a* mRNA, and channel binding leading to a covariate reduction in pain behavior.

## 1. INTRODUCTION

Human observations and pre-clinical studies have validated the role of the voltage-gated sodium channel 1.7 (NaV 1.7) as a target for pain control (1). Despite its promise as a target, there has been moderate success in creating small molecules and demonstrating efficacy in NaV 1.7-triggered pain states (2, 3).

Collective evidence suggests that NaV 1.7 transcription and expression occurs predominantly in the soma of C-polymodal nociceptors(4, 5) a conjecture which is widely accepted. Interestingly, recent evidence suggests that NaV1.7 message and protein may also be present in second and higher order dorsal horn neurons thought to participate in nociceptive processing (6, 7).

Although NaV 1.7 has a high degree of association with neuraxial nociceptive processing, its distribution has also been demonstrated in olfactory and gustatory processing systems(8), as well vagal afferent (9), and sympathetic neurons (10), raising concerns as to the potential effects of a *systemically* delivered drug on non-pain related systems. This has raised the appreciation that specifically altering expression or function of NaV 1.7 specifically in neuraxial nociceptive afferent has a particular virtue. Such anatomically selective KO has been achieved using neuraxial delivery. After such delivery, small molecules, oligonucleotides (antisense) and various viral constructs have been shown to appear in DRG neurons and to alter their transcriptional properties (11). The clinical utility of these platforms may be defined by their persistence (12).

The use of intrathecally delivered genome modifying platforms, such as viral transfection, produces a long lasting if not irreversible knock-out (KO) of protein expression, leading to persistent loss of function (11, 12). In contrast, intrathecal antisense therapeutics can in principle yield a conditional silencing of messenger ribonucleic acids (mRNAs), and long lasting, but reversible effect on protein synthesis (13–16).

In the present work, we addressed the concern that while quantification of mRNA provides an indirect index of channel expression, spatial or temporal disconnections related to protein synthesis, membrane transport, channel insertion and internalization may prevent a 1 to 1 covariance (17, 18). Here, we assessed the binding of a selective fluorophore labeled Nav 1.7 *antagonist* (ATTO488PTx-II) (19) to assess not only the amount of targeted binding at a specific site, but the fraction of cells expressing that binding. This molecule bind to the extracellularly expressed voltage sensor of the NaV1.7 complex. This approach further allows us to estimate the effect of fraction of surface expression required to produce a given effect on pain behavior (11, 12, 20). We found that antisense molecules delivered intrathecally selectively affect transcription of *Scn9a*. These molecules are taken up by DRGs, spinal cord, and brain. In addition, the dose-dependent intrathecal delivery of antisense led to a covariance of analgesia and the related ASO-evoked changes in membrane NaV1.7 binding.

## 2. METHODS

### 2.1. Animals and intrathecal drug delivery

All *in vivo* experiments were carried out according to protocols (protocol number S00137R) approved by the Institutional Animal Care Committee (IACUC) of University of California, San Diego. Male Sprague–Dawley rats (200–250 g) were prepared with lumbar intrathecal catheters as previously described by (21) and (22).

For dosing and distribution studies the Mazur et al.(22) technique was used to deliver ASO to the lumbar cord to study the ability of the NaV 1.7 antisense in knock-down *Scn9a* mRNA in dorsal root ganglia (DRGs) across the spinal column segments. The same surgical technique was used to perform the pharmacological validation and selectivity assessment from the antisense optimal dose.

After defining the optimal dose for the antisense, lumbar delivery of ASO was performed using the Yaksh technique (23) to assess the effects of *Scn9a* mRNA knock-down and reduced NaV 1.7 binding on the formalin induced flinching.

The therapeutic protocol for the delivery of ASOs started 7 days after surgery. Two antisense oligonucleotides (IONIS Pharmaceuticals Inc., Carlsbad, California, USA) were used. The NaV 1.7 ASO is an ASO that targets the *Scn9a* mRNA (NaV 1.7), where Mismatch is an ASO control. ASOs were dissolved in PBS without Calcium and Magnesium (Gibco™; 14190) and administered intrathecally as a single bolus injection (10µL) followed by a 10 or 30µL flush depending on the technique performed.

### 2.2. Assessment of nociception

In this study, three sets of nociceptive behavior were measured in rats. Initially, to evaluate the mechanical nociception two different methodologies were performed: the up-down method (24) or paw-pressure test (25). Thermal nociception was measured using a Hargreaves type device (26). Finally, the effects of IT treatments were conducted to evaluate the biphasic flinching nociceptive behavior triggered by unilateral hind paw intraplantar formalin (2.5%/50µL). To measure flinching a lightweight metal band was placed around the hind paw receiving the formalin. Movement of the band in a detection field is collected automatically and presented as flinches /min (27). In all experiments, animals were set to acclimate for at least 60 minutes prior testing.

#### 2.2.1. OD-1 flinching behavior test

OD-1 is a peptide isolated from the venom of the scorpion *Odonthobuthus doriae* which selectively modulates NaV 1.7 channels, inhibiting its fast inactivation while producing substantial and persistent propagated currents (28). The outcome is the triggering of flinching nociceptive behavior when injected into an animal’s hind paw that typically lasts up to 40 minutes (29, 30). In this study, we used OD-1 to confirm the functional knock-down of NaV 1.7 provided by the optimal dose of ASOs in rats. The flinching behavior evoked by OD-1 (500 nM; 50 μL) was analyzed in animals that were injected with the peptide through the dermal route and evaluated during a 40-minute timeframe using the automated flinch detecting system (27). Animals were set to acclimate for 60 minutes prior to the tests.

### 2.3. Defining the optimal dose of ASO

To determine the ASO optimal dose, several doses of ASOs were tested in different models of nociception. The dosing and time frame for undertaking behavioral studies was based on previous studies published by our research group and collaborators (13, 22, 31). Doses ranging from 100 to 3000µg/10μL were given intrathecally at 10-14 days prior to the tests. The animals were submitted to the assessment of nociception in formalin and OD-1 tests at day 28 after ASO delivery. In these animals, tissue collection was performed at the end of experiments. The optimal dose of ASO was chosen considering the analgesic profile observed at the behavioral tests. Once the optimal dose was selected, the analgesic profile from ASO optimal dose was evaluated in carrageenan inflammatory pain model or in paw-pressure test using a Randal-Sellito apparatus (25, 32). These two models are reported to be affected in NaV 1.7 knockouts (10).

#### 2.3.1. Selectivity assessment from antisense optimal dose

To demonstrate the selectivity profile from antisense optimal dose, animals were previously treated with NaV 1.7 ASO, Mismatch or phosphate buffered saline (PBS) controls. On day 14 after injections, these animals were euthanized and L3-L5 DRGs were collected and processed for quantitative mRNA assessment from several NaV subunits: *Scn1a* (NaV1.1), *Scn2a* (NaV 1.2), *Scn8a* (NaV 1.6), *Scn9a* (NaV 1.7), *Scn10a* (NaV 1.8) and *Scn11a* (NaV 1.9). All these channels are reported to be expressed in DRG sensory neurons and contribute to pain signaling (33–35).

#### 2.3.2. Tolerability assays

Tolerability was assessed for the optimal dose of NaV 1.7 ASO by: I) immunohistochemistry (IHC) and quantitative measuring of mRNA for markers of glial activation: *Aif-1* (Ionized Calcium Binding Adaptor Molecule 1 -IBA-1) and *Gfap* (Glial fibrillary acidic protein-GFAP). The mRNA levels and/or IHC outcomes from these markers were measured in thoracic DRGs and spinal cords, and at the cerebral cortex harvested from NaV 1.7 ASO, Mismatch, and PBS injected animals.

##### 2.3.2.1. Histopathology

We used immunohistochemistry to perform histopathology assays and analyze glial reactivity in DRGs, spinal cords, and Brain through IBA-1 staining. Gliosis was determined following the score system in Table 2 where activation was taken as an indicator of treatment related toxicity (36).

##### 2.3.2.2. Antisense uptake studies

A second set of experiments was conducted in DRGs, Spinal cords, and Brain, to analyze the ASO uptake. Here, the tissue was fixed and then processed on a mouse brain protocol using a Sakura Tissue Tek tissue processor. After embedding, slides were cut at 4 microns, air dried overnight then dried at 60° for 1 hour. Slides were stained with Rabbit polyclonal ASO (IONIS) antibody on a Ventana Ultra staining system. ASO slides were treated enzymatically with Trypsin (Sigma, T8003). The slides were then blocked with Endogenous Biotin Blocking Kit (Ventana, 760-050) and Normal Goat Serum (Jackson Immuno Labs, 005-000-121). The primary antibody was diluted with Discovery Antibody Diluent (Ventana, 760-108) and incubated for 1 hour at 37°c. The antibodies were detected with Biotin labeled Goat Anti-Rabbit secondary antibody (Jackson Immuno Labs, 111-005-003). The secondary antibody was labeled with DABMap Kit (Ventana, 760-124). The Iba1 slides were treated with Heat Induced Antigen Retrieval (HIER) with Ventana CC1 solution (Ventana, 950-500) for 64 minutes. The primary antibody was diluted with Discovery Antibody Diluent (Ventana, 760-108) and incubated for 1 hour at 37°c. The antibody was detected with UltraMap anti Rabbit Polymer detection (Ventana, 760-4315). The rabbit polymer was labeled with Ventana ChromoMap DAB kit (Ventana, 760-159). Images were scanned on a Hamamatsu S360 scanner at 20X resolution. The histopathological scoring and ASO uptake were analyzed by an investigator without knowledge of treatment.

### 2.4. In vitro characterization of ATTO488PTx-II binding in Nav1.7 transfected HEK cells

To analyze the feasibility of the ligand binding approach, we initially constructed a model system using Human Embryonic Kidney (HEK) cells transfected with HaloTag-NaV 1.7 DNA sequences. In this system, the NaV 1.7 channels can be labeled using a specific TMR ligand that targets the Halo-tag expressed at NaV 1.7 proteins (37, 38). The purpose was to define the optimal concentration to label NaV 1.7 channels using the ATTO488-PTx II antagonist (488-PTx II), a fluorescent NaV 1.7 antagonist (19). Briefly, cells were plated onto glass coverslips coated with poly-D-lysine at 20-30% confluence. After 24 hours, half of the wells of a 12-well plate were transfected using 1 μg of plasmid carrying a mouse NaV 1.7-HaloTag DNA sequences, along with 20 μL of Lipofectamine 2000 (transfection reagent). The plasmid was kindly provided by Dr. Rajesh Khanna (University of Florida, Gainesville, Florida, USA). The media was replaced 4 hours after the start of the transfection. After 24 hours, cells were placed on ice and stained with a Halo ligand (Promega®, G825A, cell permeable TMR ligand; 2.5 μM final concentration) and ATTO488PTx-II (Smartox®; 0.5, 2.5 or 10 μM final concentration) for 30 minutes. After one wash in Tyrode solution (as recommend by (19), cells were fixed in 4% PFA and mounted onto microscope slides. Each concentration of ATTO488PTx-II was examined in 3 transfected samples and 1 untransfected sample (used as a control for nonspecific binding). Samples were imaged with a Leica TCS SP5 confocal system to determine the fluorescence intensity, covariance, and colocalization between the signal of two ligands. In all images obtained, the data was analyzed using the LASX Office version 1.4.7 software.

### 2.5. Dorsal root ganglia cell culture

Immediately after the behavioral tests, rats were euthanized under deep anesthesia using 5% Isoflurane followed by decapitation. The DRGs were harvested and underwent dissociation to perform cell culture as previously described (39) with some modifications. The Ganglia were incubated with 2% collagenase (type IV; Gibco®; catalog number 17104019) for 120 minutes and 0.25% Trypsin (Gibco®; catalog number 25200056) for 15 minutes. After two washes with DMEM (Gibco®), the ganglionic cells were dissociated using glass Pasteur pipettes followed by filtration through an 70µm cell strainer for debris removal, and centrifugation at 1400 rpm for 10 minutes. The cell pellet was resuspended in DMEM and subsequently submitted to 10% bovine serum albumin gradient separation followed by centrifugation at 1400 rpm for 10 minutes. The residual debris were removed and the cells plated in 25 mm coverslips coated with poly-D-lysine/Laminin (0.02mg/mL; 0.016 mg/mL, respectively) and further disposed on 35mm single-dish cell culture plates (Thermo scientific^TM^; catalog number 130180). All cultures were maintained in DMEM plus 1% penicillin (50 U/mL)/streptomycin (50 mg/mL), 10% Fetal bovine serum (Sigma-Aldrich; catalog number F2442), at 37 °C with 5% CO_2_ until the beginning of experiments or mRNA extraction.

#### 2.5.1. Tissue processing for binding assays

For binding assays, the ipsilateral or contralateral L1-L5 DRGs from n = 4-10 animals were pooled and used to obtain DRG cell cultures as previously described. The cell pellets were divided into 2 pools. The 1^st^ pool consisted of 1×10^4^ cells (6 wells per group in each set of experiments) plated in a 96-well plate coated with poly-D-lysine/Laminin. After a 24-hour plating, these cells were lysed with RLT Buffer (QIAGEN®) and further analyzed for *Scn9a* mRNA quantification through qPCR. The 2^nd^ pool consisted of cells plated in 25 mm poly-D-lysine/Laminin coated coverslips at 40-70% confluence. After 24-hour plating, these cells were employed in binding assays with ATTO488PTx-II and further fluorescence imaging.

### 2.6. Imaging experiments

#### 2.6.1. Assessment of ATTO488PTx-II binding in dorsal root ganglia cell cultures

To determine the NaV 1.7 selective binding in under naïve conditions in NaV 1.7 ASO and Mismatch groups, rats were injected with the optimal dose of NaV 1.7 ASO and Mismatch and sacrificed at day 14^th^ after ASO delivery. The L1-L5 DRGs were bilaterally harvested to obtain DRG cell cultures as previously mentioned (item 2.5). The binding for ATTO488PTx-II (488-PTxII) was assessed in 24h-plated DRG cell cultures and performed as follows: cells were washed once with Tyrode solution for 5 minutes following blockade with 2% BSA for 30 minutes and staining with 10 μM ATTO488-PTx II (diluted in Tyrode solution) for 30 minutes. Next, coverslips were fixed with 4% PFA and the slides were mounted using prolong gold (Invitrogen®) containing DAPI. All 488-PTx II binding assays were carried out on the same day and the coverslips were maintained on ice to minimize channel endocytosis (37). Controls consisting of cells treated with Tyrode solution without 488-PTx II were employed to assess non-specific binding. Two sets of binding analysis were performed: **i)** different slides were analyzed and an average of 70 cells expressing fluorophore were counted in each group. The normalized cell fluorescence was calculated by the following formula: (Integrated Density of cell fluorescence) – (Average Mean of Background fluorescence intensity x Cell area in micrometers); **ii)** We determined the relative percentage of cells in these analyzed slides aiming to see the remaining fraction of cells still expressing NaV 1.7 signal according to soma size. A range between 6-11 fields were analyzed in each group and the relative percentage of NaV 1.7 positive cells were calculated by the formula: [(Number of ATTO488-PTx II positive cells **÷** Number of DAPI positive cells)] x 100. A minimum of 10 cells were required in each field, and only cells displaying clear visibility from the membrane perimeter were analyzed.

The basal levels of ATTO488PTx-II fluorescence in NaV 1.7 ASO and Mismatch controls were determined using an Olympus BX51 fluorescence microscope. The coverslips were mounted in microscope slides with prolong gold containing DAPI and imaged at 63x magnification. In this set of experiments, the data was analyzed using the Image J software.

#### 2.6.2. Assessment of ATTO488PTx-II binding in dorsal root ganglia cell cultures obtained from ASO -treated rats submitted to the formalin test

At the time frame of 24 hours after the formalin test, Lumbar (L1-L5) DRGs cell cultures (one from each anatomic side: ipsilateral or contralateral) from NaV 1.7 ASO (300 and 1000 µg/10µL doses) and Mismatch (1000 µg/10µL) groups were obtained following the protocol previously described on item **2.5**. As described on item **2.5.1**, the cell culture pools were plated differently for NaV 1.7 binding analysis and *Scn9a* mRNA quantification. After 24 hours, cultures corresponding to each treatment set were stained with ATTO488PTx-II and lysate for mRNA extraction as described on item **2.5.1**. The intensity of fluorescence in each group was assessed in whole-cell z-stacks and assumed as indicative from amount of NaV 1.7 binding calculated as previously described on item **2.6.1.**

#### 2.6.3. Z-stacks of dorsal root ganglia cell cultures

We used a Leica TCS SP5 confocal system to determine the fluorescence intensity of ATTO488PTx-II signal in z-stacks acquired from DRGs cell cultures obtained from NaV 1.7 ASO and Mismatch groups that were harvested after the formalin test. To assess the ligand signal across the z-axis in the cells, the system was set-up to perform readings at 63x magnification, using a 0.1µm step-size and at 400Hz speed. In each slide imaged, an average of 5 fields were recorded and the intensity of ligand fluorescence measured in the max projection images of z-stacks using the Leica imaging software LASX Office version 1.4.7.

#### 2.6.4. Assessment of ATTO488PTx-II binding in sciatic nerves obtained from animals submitted to the formalin test

At the time frame of 24 hours after the formalin test, the sciatic nerves from each animal of NaV 1.7 ASO (300 and 1000 µg/10µL doses) and Mismatch (1000 µg/10µL) groups were collected. The proximal section of the nerve was frozen on dry ice and further addressed for *Scn9a* mRNA quantification. The medial section of the sciatic nerve was placed on 4%PFA for fixation. After 24hs, the nerves were gradually embedded on 10-30% sucrose solution and finally placed on O.C.T compound (Tissue Tek, Sakura ^TM^, Japan) cryo-solution to be cryo-sectioned at −20°C in a cryostat. Sections of 20 μm from ipsilateral sciatic nerves were obtained and placed on slides (Superfrost plus; Fisherdbrand®) to be stained with ATTO488PTx-II. Slides were dried at room temperature for 30 minutes, followed by a series of washes with 0.01M PBS (pH 7.4). Unspecific binding of the secondary antibody was prevented by incubating the tissue with a blocking solution containing 3% normal donkey serum (Millipore^TM^, S30), 0.3% Triton X-100 (Sigma, cat: 9036-19-5) in 0.01M PBS for 60 minutes. Thereafter, slides were incubated at 4°C overnight with an anti-PGP9.5 antibody (1:2000, Cedarlane, CL7756) or during 48 hours with ATTO488PTx-II. After the incubation with PGP9.5, slides were incubated with a donkey anti rabbit Cy2 antibody (1:200, Jackson ImmunoResearch, 711-005-152) for 2 hours, followed by a conterstainig with DAPI (1:1000) and coverslipped with ProLong Gold antifade Mounting (Invitrogen, P36934). Slides incubated with ATTO488PTx-II, were followed by counterstaining with DAPI and coverslipped with ProLong Gold antifade mounting. Slides were kept at room temperature for 24 hours before imaging. Z-stack images were acquired using a Zeiss LSM 800 microscope. Images were transformed into maximum projections with ZEN software from Zeiss. A single ROI was acquired from five different sciatic nerve sections separated by 40 μm each.

### 2.7. RNA extraction and quantitative real-time PCR analysis

At the completion of an *in vivo* study, animals were deeply anesthetized with isoflurane and tissue harvesting was performed according to each study experimental design. For analysis of DRG neuronal cell cultures, cells were dissociated and plated. After 24 hours cells were washed three times for 5 minutes with Dulbecco’s phosphate buffered saline (dPBS 1x; Gibco®) and then lysate with RLT buffer (Qiagen®) at 100 μL/well for RNA extraction following manufacturer’s instruction [Qiagen, Valencia, CA]. For tissues, samples were homogenized in of Trizol reagent (Thermofisher scientific, Waltham, MA) and total RNA was extracted using the Life Technologies mini-RNA purification kit (Qiagen, Valencia, CA) according to the manufacturer’s protocol. After purification, the RNA samples were subjected to real-time RT-PCR analysis using the Life Technologies ABI QuantStudio 7 Flex Sequence Detection System (Applied Biosystems Inc, Carlsbad, CA). Briefly, 10 µl RT-PCR reactions containing 400 nl of RNA were run with the AgPath-ID One-Step qRT-PCR Kit (Thermofisher scientific, Waltham, MA) reagents and the primer probe sets (Table-1). All qPCR reactions were run in triplicate. Outliers in triplicates due to technical issues were eliminated. Target mRNA was then normalized to Ppia, a ubiquitously expressed housekeeping gene, and this was further normalized to the level measured in control animals that were administered PBS.

**Table-1.**
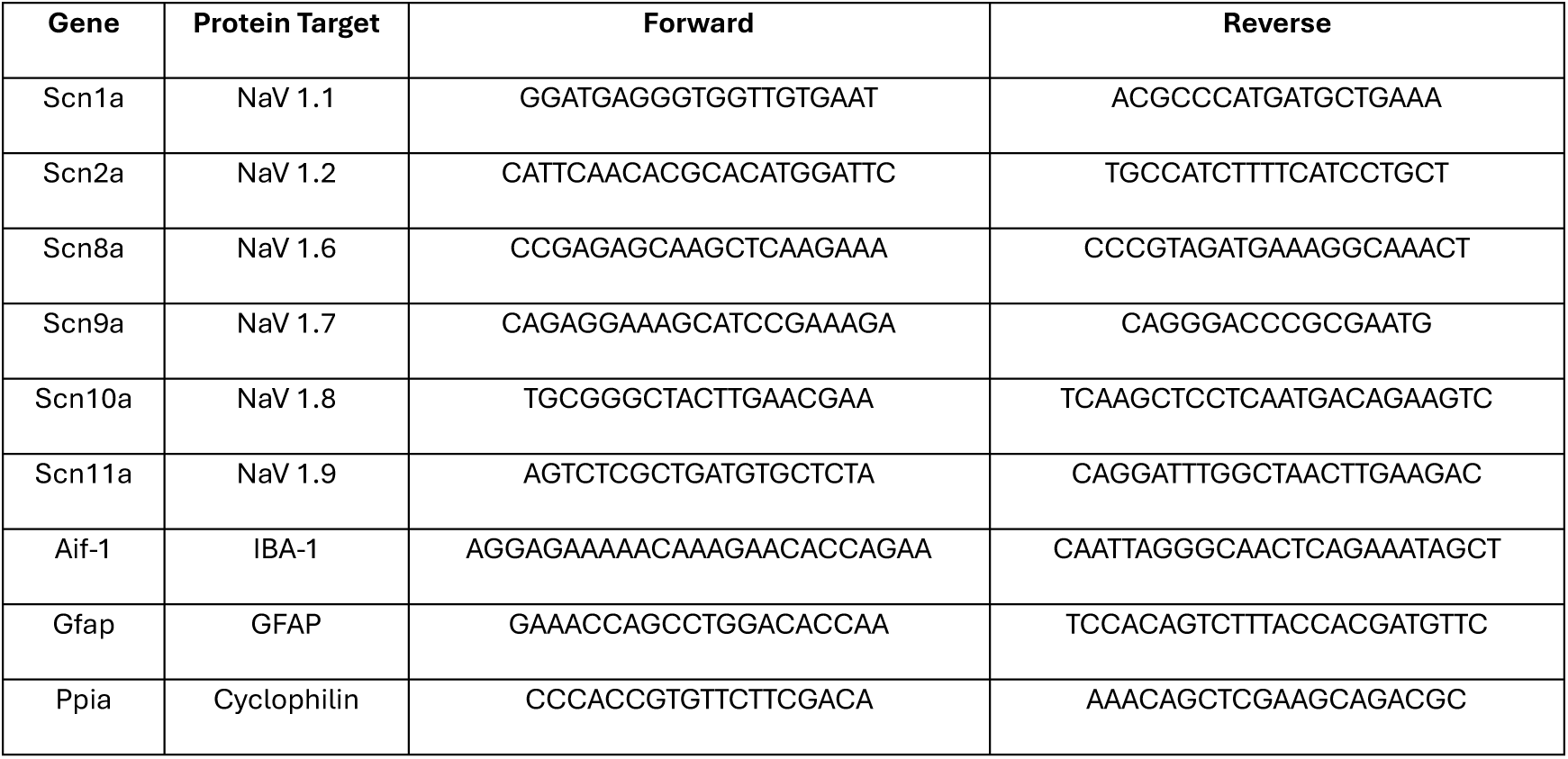
Primers used in this study.

**Table-2.**
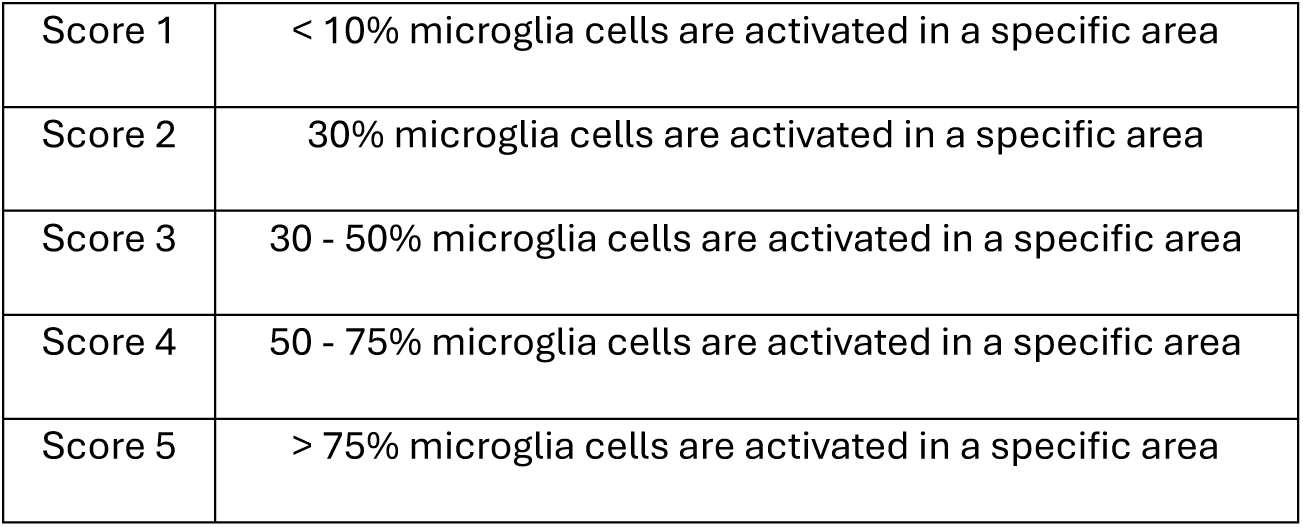
Score system used to assess microgliosis across dorsal root ganglia, spinal cord, and brain.

### 2.8. Statistical analysis

The analysis of the results was performed using GraphPad Prism v.10 software (GraphPad®, San Diego, USA). The normal distribution of values in each analysis was performed with the D’Agostino Pearson, Shapiro-Wilk or Kolmorogov-Smirnov test, according to the sample size. When a non-parametric statistical test was performed, the Mann-Whitney test was used to compare two means. When the comparison involved more than two means, the Kruskal-Wallis test was performed with Dunn’s post-hoc test to determine if there were differences between the groups analyzed. If the values followed a normal distribution, parametric statistical tests were performed. The T-student test was selected when two means were compared. When the comparison involved more than two means, a one-way or two-way analysis of variance (ANOVA) was performed according to the experimental design followed. When the level of significance indicated a statistical difference between the analyzed means, the Tukey or Dunnet’s test was used to compare the different analyzed groups. The level of significance adopted in each analysis of the results was p < 0.05.

## 3. RESULTS

### 3.1. Determining the optimal dose of ASO – Distribution studies

The injection of different concentrations of ASOs evoked a caudal to rostral gradient of knockdown of *Scn9a* mRNA in DRGs. At lumbar levels, significant knockdown vs missense was achieved with the 100 μg dose, where effective knockdown at cervical levels was only possible with 1000 and 3000 μg doses (Figure 1A-C). These highest doses displayed similar outcomes: decreasing in 50% the *Scn9a* mRNA in comparison with PBS controls. As no differences were seen between the highest doses (1000 and 3000 μg) regarding knockdown activity, we assumed 1000 μg as being the minimum optimally effective ASO dose. Given the broad DRG-distribution across the spinal column, we investigated the effects of this optimal dose in trigeminal ganglia, as well as evaluated its selectivity to address NaV 1.7 specific knockdown at lumbar DRGs. The ASO optimal dose led to a significant knockdown of *Scn9a* mRNA at the trigeminal ganglia (Figure 1D) In these studies we further examined the selectivity of the active ASO NaV 1.7 (Figure 1E) In these studies no significant reduction was observed for NaV 1.7 antisense in the mRNA related to other NaV channel subunits (NaV 1.1-NaV 1.9).

**Figure 1.**
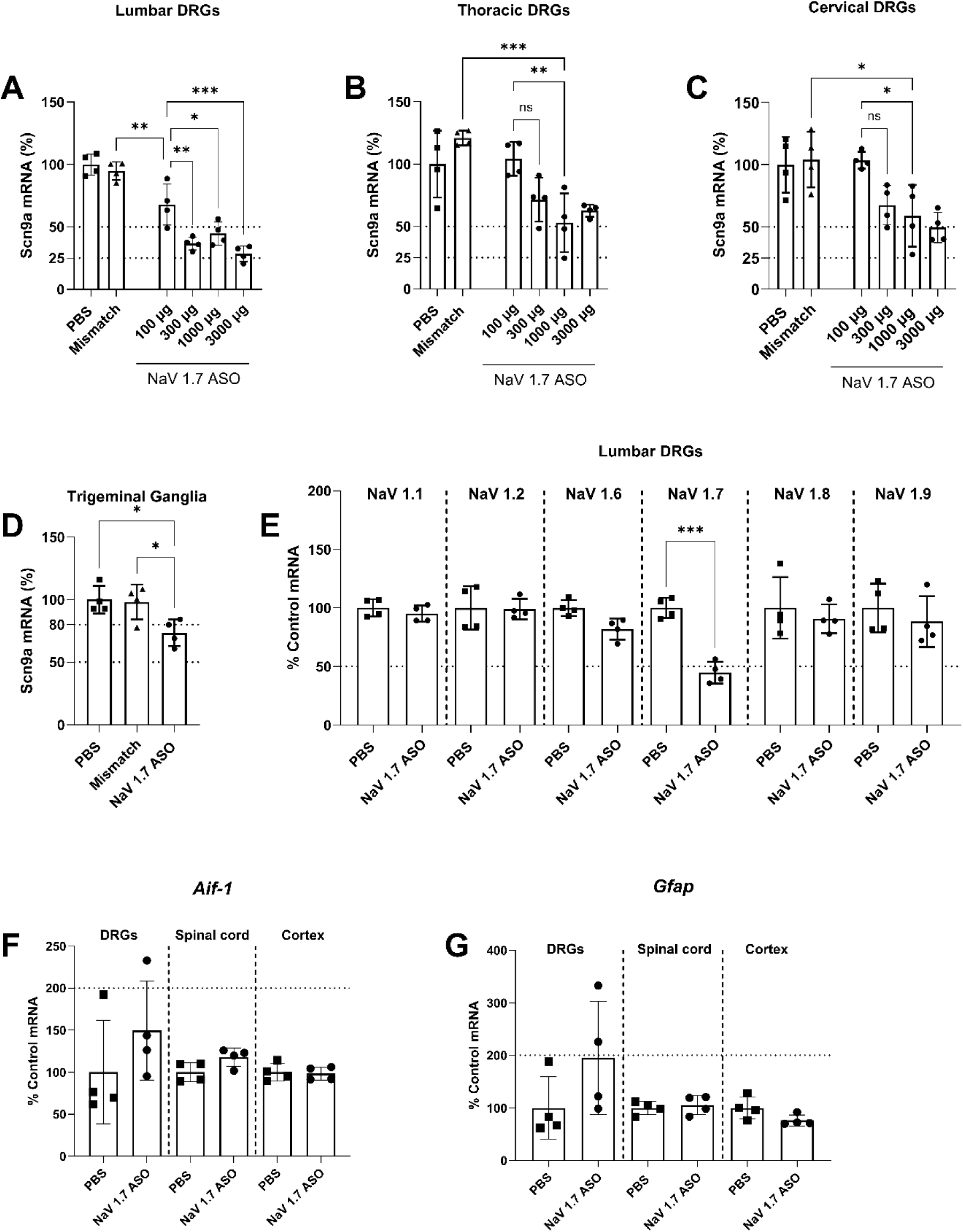
Dose-dependent knock-down of *Scn9a* mRNA in sensory ganglia across the spinal cord axis. The intrathecal injection of NaV 1.7 ASO (100/300/1000 or 3000 μg; 10 μL) dose-dependently induces a knock-down of *Scn9a* in lumbar, thoracic and cervical dorsal root ganglia (A-C). The effective dose of NaV 1.7 ASO (1000 μg; 10 μL) significantly knock-down *Scn9a* mRNA in trigeminal ganglia (D) and selectively affects NaV 1.7 mRNA at the lumbar DRGs without interference in other voltage-gate sodium channels subtypes (E). On (F-G) Assessment of DRGs, spinal cord and brain cortex toxicity related to the optimal dose of NaV 1.7 ASO (1000 μg; 10 μL). This dose does not lead to significant transcriptional alterations in glial activity markers *Aif-1* (F-IBA-1 protein, macrophages/microglia) and *Gfap* (G-GFAP protein, satellite glial cells/astrocytes) in DRGs, spinal cord and Cortex samples of ASO animals in comparison with PBS controls. Results shown as mean ± standard deviation. Symbols *, ** and *** indicate P<0.05, P<0.01 or P<0.001 for comparison between groups. On (A-D) One-way ANOVA with Tukey post-hoc test. On (E, F, and G) T-student test.

### 3.2. Assessing the ASO uptake profile for the optimal dose

#### 3.2.1. Dorsal root ganglia

In dorsal root ganglia harvested on the 14^th^ day after ASO delivery, ASO particles of both mismatch and NaV 1.7 antisense could be detected incorporated by large and small neurons (Figure 2A-D; E-H; Black and blue arrows, respectively). Antisense particles could also be detected at nerve filaments projecting across the DRG (Figure 2A, 2G, asterisks) for both treatment groups.

**Figure 2.**
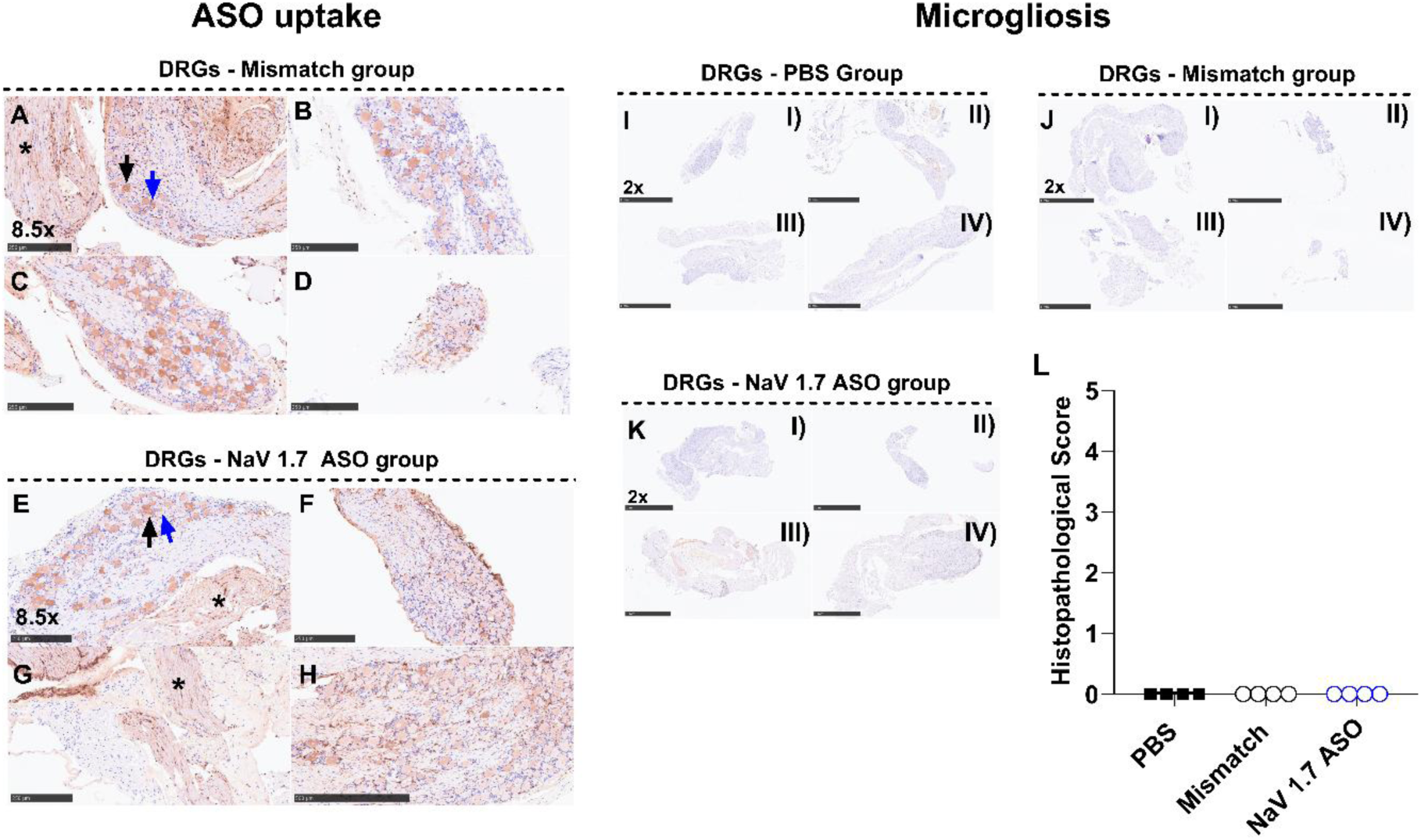
Uptake of oligonucleotides by dorsal root ganglia neurons and associated microgliosis. The lumbar intrathecal delivery of Mismatch or NaV 1.7 antisense lead to rostral distribution and antisense uptake at thoracic dorsal root ganglia. Both Mismatch (A-D) and NaV 1.7 (E-H) antisense particles can be seen incorporated by large (black arrows) and small (blue arrows) neurons. In both groups, a residual staining for antisense particles can be seen at nerve filaments projecting across the DRG (A, G, asterisks). The intrathecal injection of NaV 1.7 ASO optimal dose (1000 μg; 10 μL) does not lead to an increase of IBA-1 staining (microgliosis) in comparison with Mismatch and PBS controls (I, J, or K, I-IV). No tissue injury was observed between DRGs harvested from Mismatch, NaV 1.7 ASO in comparison with samples from PBS control group, therefore no significant changes in the histopathological score were observed between the groups analyzed (L). Results shown as mean ± standard deviation. Kruskal-Wallis test.

#### 3.2.2. *Spinal Cord* and Brain

In spinal cords harvested on the 14^th^ day after ASO delivery, ASO particles of both mismatch and NaV 1.7 antisense could be detected incorporated by cells of superficial and deep dorsal horn (Supplementary figure 2A and C). In brains harvested on the 14^th^ day after ASO delivery, ASO particles of both mismatch and NaV 1.7 antisense could be observed to be moderately incorporated by cells throughout the whole brain (Supplementary figure 2B and D).

### 3.3. Assessing the toxicity of ASO at DRGs, spinal cords, and brain

In dorsal root ganglia harvested on the 14^th^ day after ASO delivery, the NaV 1.7 antisense optimal dose induced a slight increase in the mRNA levels of glial markers *Aif-1* (IBA-1) and *Gfap* (GFAP), however, not significantly different from PBS controls (Figure 1F-G). In addition, no significant changes in these markers were detected spinal cords and cortex harvested from the same groups (Figure 1F-G).

At the histopathological analysis, mismatch and the NaV 1.7 antisense optimal dose intrathecal delivery induced a slight gliosis at spinal cord, and brain, however, the score findings were not different from PBS controls (supplemental figure 3 AI-VI, B, and C).

### 3.4. ASO targeting NaV 1.7 dose-dependently affects nociceptive behavior evoked by formalin

The intrathecal injections from a series of NaV 1.7 antisense doses (100-3000 μg; 10 μL) significantly knockdown *Scn9a* mRNA in DRGs and decreased formalin second phase flinching (Figure 3A and B). Interestingly, despite the lower doses of ASOs, 100 and 300 μg/10μL provided robust knock-down of NaV 1.7 mRNA, significant effect on formalin nociceptive response were only seen for rats treated with 1000 and 3000 μg doses (Figure 3A-B). Although no further degree of knock-down was obtained with doses greater than 1000 μg, rats treated with NaV 1.7 ASO at 3000 μg/10μL had an even lower nociceptive response in comparison with animals treated with the 1000 μg dose (Figure 3A).

**Figure 3.**
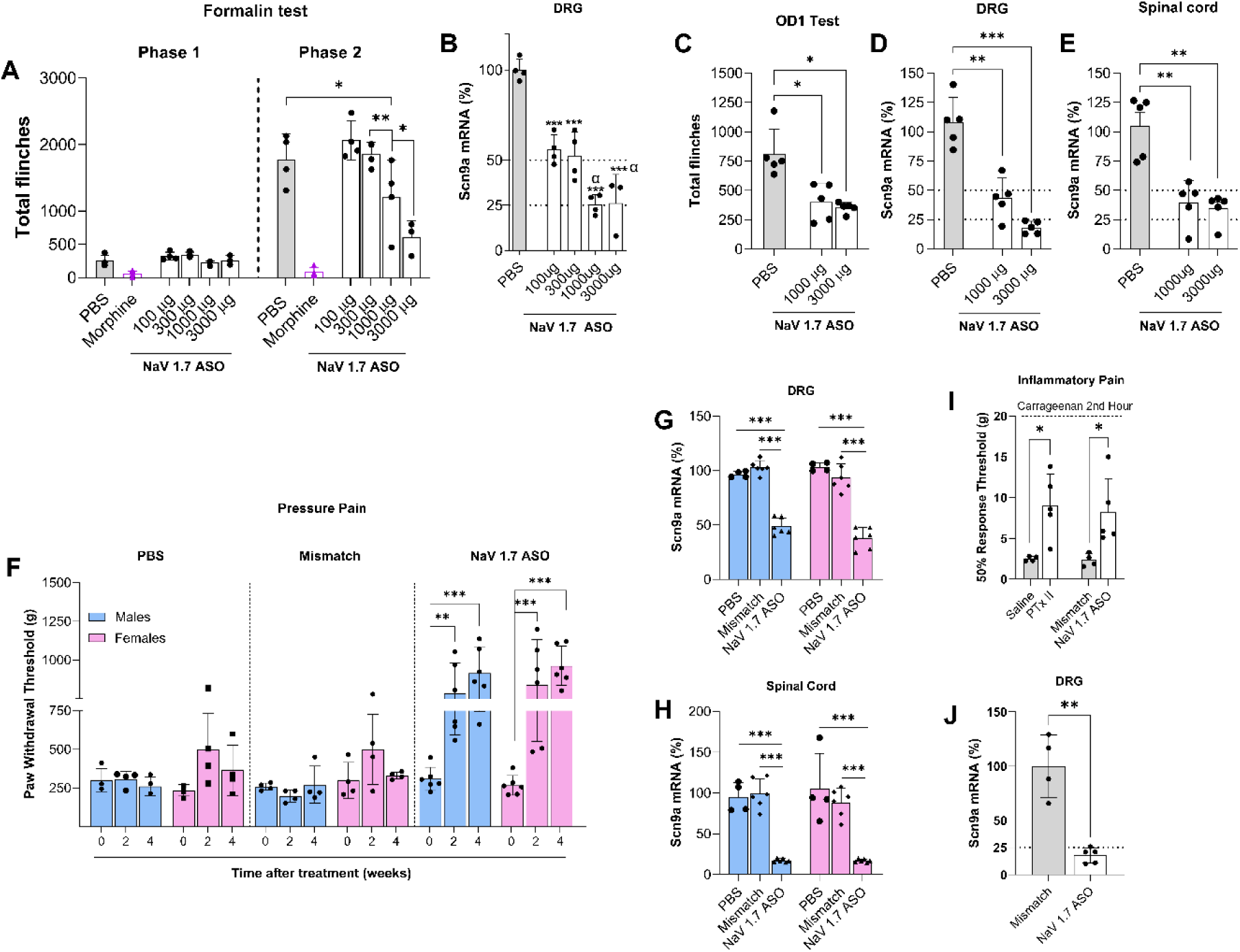
ASOs targeting NaV 1.7 affects nociceptive behaviors. The intrathecal injections of different NaV 1.7 ASO doses (100-3000 μg; 10 μL) significantly knock-down *Scn9a* mRNA in DRGs and decreased formalin second phase (**A, B**). The nociceptive behavior evoked by the selective NaV 1.7 agonist, OD-1, was significantly affected by the intrathecal knock-down of NaV 1.7 with 1000 and 3000 μg ASO dose **(C)**. These doses tested had similar outcomes in decreasing the flinching behavior triggered by intradermal injection of OD-1 with similar degrees in message knock-down at the DRG **(D)** and spinal cord **(E)**. On a 4-week time course, the intrathecal NaV 1.7 antisense markedly increase the paw withdrawal thresholds on males and females at week 2 and 4 after the ASO delivery (**F**). These events are correlated with a knock-down of *Scn9a* at both DRGs (**G**) and spinal cord (**H**) in comparison with Mismatch and PBS controls. In an extra-set of experiments (**I**) the intrathecal NaV 1.7 ASO (1000 µg; 10 µL; intrathecal; 14 days prior testing) replicates the analgesic profile of the NaV 1.7 antagonist PTx-II (2 µg; 10 µL; intrathecal) in the carrageenan model (100 µg; 50 µL; subcutaneously; right-hind paw). Again, the effects of NaV 1.7 ASO correlates with a significant knock-down in *Scn9a* mRNA in the ipsilateral L3-L5 DRGs (**J**) harvested from mismatch and NaV 1.7 ASO right after testing. Results shown as mean ± standard deviation. On **(A)** symbols * and *** indicate P<0.05 and P<0.001 for comparison with PBS, while symbols # and & indicates P<0.05 for comparison respectively with 300 μg and 1000 ASO doses. On **(B)** symbols * and # indicates P<0.05 for comparison with PBS or 300 μg ASO dose. On **(C)**, **(D)**, and **(E)** symbols *, **, and *** indicate P<0.05, P<0.01, or P<0.001 for comparison between groups. Two-way ANOVA with Tukey post-hoc test on **(A)**. On (**B-E**) One-way ANOVA with Tukey post-hoc test.

### 3.5. ASOs targeting NaV 1.7 decrease nociceptive behavior evoked by OD1

After investigating the effects of NaV 1.7 knockdown in formalin model, we evaluated the capability of ASOs in modulating the intraplantar OD-1 flinching response, a NaV 1.7 agonist. For this experiment, we selected the 1000 and 3000 μg dose of ASO given their ability in providing significant *Scn9a* mRNA knock-down and affect nociceptive response at the formalin test. As shown in figure 3C-E, the nociceptive behavior evoked by the selective NaV 1.7 agonist, OD-1, was significantly affected by the intrathecal knockdown of *Scn9a* mRNA. Both ASO doses tested had similar outcomes in decreasing the flinching behavior triggered by intradermal injection of OD-1 (Figure 3C) with similar degrees in message knockdown at the DRG (Figure 3D) and spinal cord (Figure 3E). These results suggest that intrathecal knockdown of NaV 1.7 towards silencing mRNA via ASO injection led to a decrease in nociceptive behavior triggered by selective activation of NaV 1.7. Such analgesic effect is probably due to the silencing of *Scn9a* mRNA and further decreased expression of NaV 1.7 channel along the cell body and terminals of primary afferent neurons, affecting the nociceptive signal conductance triggered by NaV 1.7 activation. As the 1000 μg and 3000 μg ASO doses showed similar outcomes in mRNA knockdown and antinociceptive response, the 1000 μg/10μL was selected as ASO optimal dose and was used in further experiments.

### 3.6. ASO targeting NaV 1.7 provides analgesia in males and female rats

After a series of behavioral experiments to set the ASO optimal dose, we investigated the analgesic activity of the optimal dose from NaV 1.7 ASO in males and female rats. For this purpose, we evaluated the paw withdrawal thresholds for nociceptive response evoked by deep pressure through the randall-sellito test in males and female rats subjected to NaV 1.7 knockdown. As shown in figure 3F-H, the selective knockdown of NaV 1.7 at DRGs and spinal cords (Figure 3G-H) markedly affected the mechanical nociception of NaV 1.7 ASO male and female rats, that developed significant increases in paw withdrawal thresholds at 2 and 4 weeks after ASO intrathecal delivery (Figure 3F). This increase in mechanical nociceptive thresholds was not seen in Mismatch and PBS controls (Figure 3F).

### 3.7. ASO targeting NaV 1.7 provides analgesia in inflammatory pain model

After confirmed the analgesic effect from the NaV 1.7 antisense in male and female rats, we investigated the ability of ASO optimal dose in provide analgesia in a model where NaV 1.7 blockers as protoxin-II(40) would induce an analgesic profile, as in carrageenan inflammatory pain model (41). As shown in figure 2I, both NaV 1.7 antagonist or antisense given intrathecally, were able to reverse the tactile allodynia in comparison with controls, saline or mismatch. The effects of NaV 1.7 ASO are attributed to an effective knockdown of *Scn9a* mRNA at the DRGs (Figure 3J) in comparison to mismatch. The analgesic profile evoked by both NaV 1.7 antagonist and antisense at the same model highlights the covariance between transcriptional alterations and target functionality.

### 3.8. Assessing the binding of ATTO488PTx-II in HEK cell platform

We transfected the NaV 17-Halotag channels at the HEK cell platforms used in this study (Supplementary figure 1A, TMR-Halo tag signal). The HEK cells transfected with the NaV1.7-HaloTag plasmid exhibited significant fluorescence signals in comparison with untransfected controls (Supplementary figure 1C). As previously mentioned, several doses were tested on our protocol to assess the ATTO488PTx-II binding in NaV1.7-Transfected HEK cell platform. Only the 10 μM dose provided a significant signal in comparison with controls (Supplemental Figure 1A-C). This fluorescence generated by the antagonist binding displayed strong covariance with the Halo-tag fluorescence, indicating colocalization with the NaV 1.7 expressed channels (Figure 4B-C). In our hands, the ATTO488PTx-II was able to generate a colocalized signal of 60% with the NaV 1.7-Halotag channels expressed (Figure 4DI-IV, and 4E-G), as shown on the Pearson’s correlation coefficient (r) (0.6440 ± 0.0841) and in the Mander’s coefficient (0.6631 ± 0.0768).

**Figure 4.**
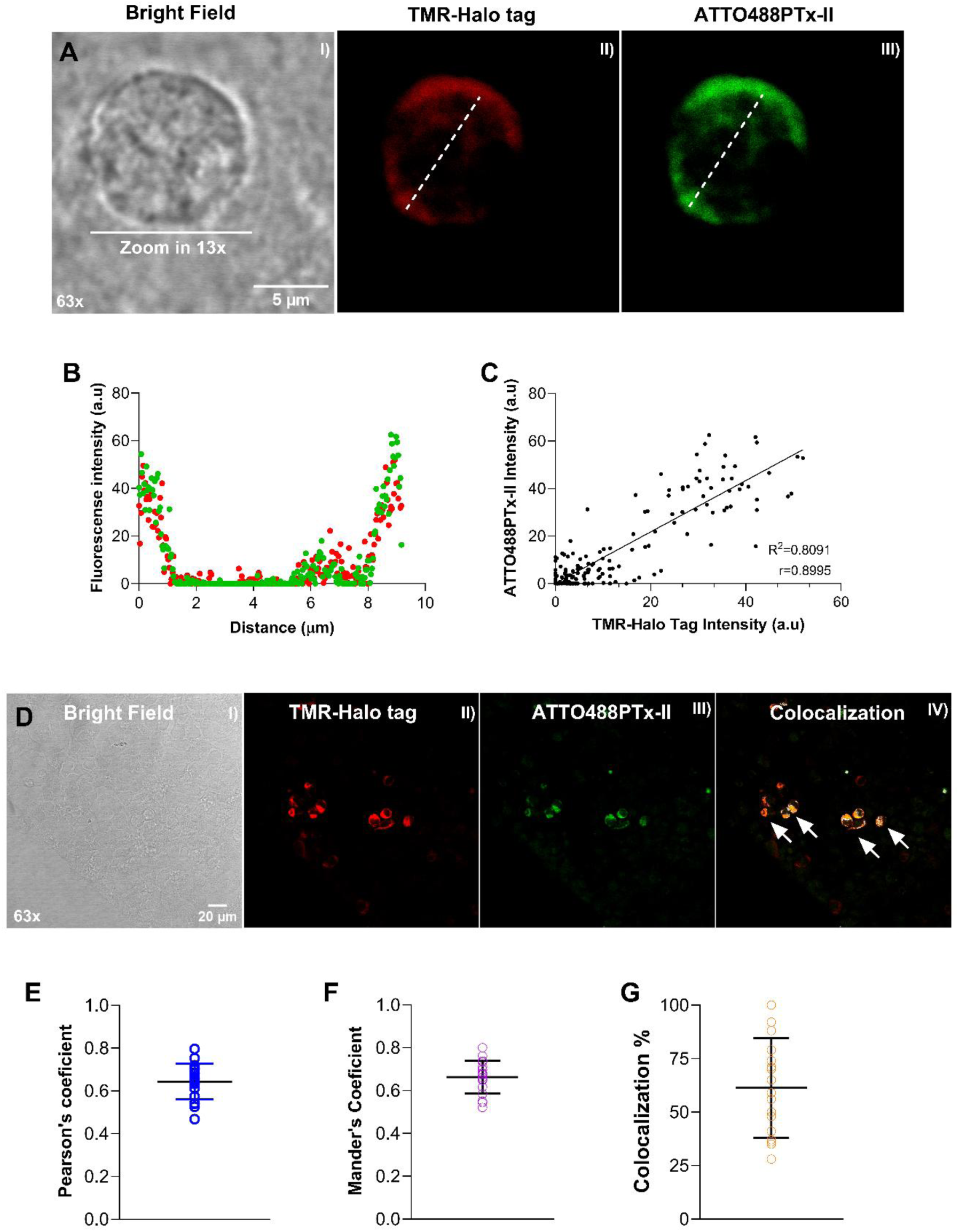
The ATTO488PTx-II signals colocalizes with TMR-ligand dye indicating a selective binding with the NaV 1.7 antagonist. **(A, I-III) (B)** Signal covariance analysis performed showing the fluorescent profile plot from the cell transfected with NaV 1.7 plasmid and stained with TMR-Ligand + 488-PTxII. The corresponding correlation analysis between the signal intensity for the two ligands is shown in **(C)**. The HEK cells expressing NaV 1.7 exhibited colocalized dots shared between the stained sites for 488-PTx II and TMR ligand (**DI-IV**) with an average of 60% colocalization **(G)**. Overall, both Pearson’s and Mander’s correlations in all samples analyzed highlighted the colocalization data showing an average correlation **(E)** and an overlapping coefficient **(F)** around 0.6. In **(B)** results are shown as the mean fluorescence intensity detected for both ligands (Y axis) across the distance analyzed (x axis). In **(C)** results shown as the mean fluorescence intensity detected (Y axis 488-PTx II; X axis TMR-ligand for Halo tag) for the ligands. In **(E)**, **(F)**, and **(G)** the results are present as mean ± standard deviation and shows, respectively, the data acquired for pearson’s and mander’s coefficient, and colocalization.

### 3.9. ATTO488PTx-II binding is reduced by Nav 1.7 antisense at small DRG neurons

After pharmacologically validating the NaV 1.7 ASO and confirming the feasibility of NaV 1.7 binding approach at the HEK cell platform, we next conducted a series of experiments to elucidate the binding phenotype in animals treated with NaV 1.7 antisense (A brief experimental design is displayed in figure 5D). Initially, our approach to label NaV 1.7 channels was replicated in Naïve DRG cell cultures (Figure 5A, B, I-III). The DRG neurons labeled with 488-PTx II exhibited visible fluorescence signals in comparison with non-labeled controls (Figure 5A, B, I-III). Some neurons displayed punctate signal in the axon terminals and towards the cell body (Figure 5C I-IV), which was confirmed through densitometry analysis (Figure 5V). The intrathecal NaV 1.7 antisense led to a visible reduction of 488-PTx II signal in DRG cells in comparison to mismatch controls (Figure 5E, I-VI). The quantitative fluorescence intensity for 488-PTx II in DRG cells of NaV 1.7 ASO animals was dramatically reduced in comparison with mismatch controls (Figure 5F). The majority percentage of cells expressing NaV 1.7 binding for 488-PTx II are small-diameter neurons (Figure 5G-I). This percentage is reduced in NaV 1.7 ASO animals suggesting that the ASO treatment was able to knock down the NaV 1.7 mRNA and lead to an absent binding for the NaV 1.7 antagonist due to lack of expression of the channel in some cells.

**Figure 5.**
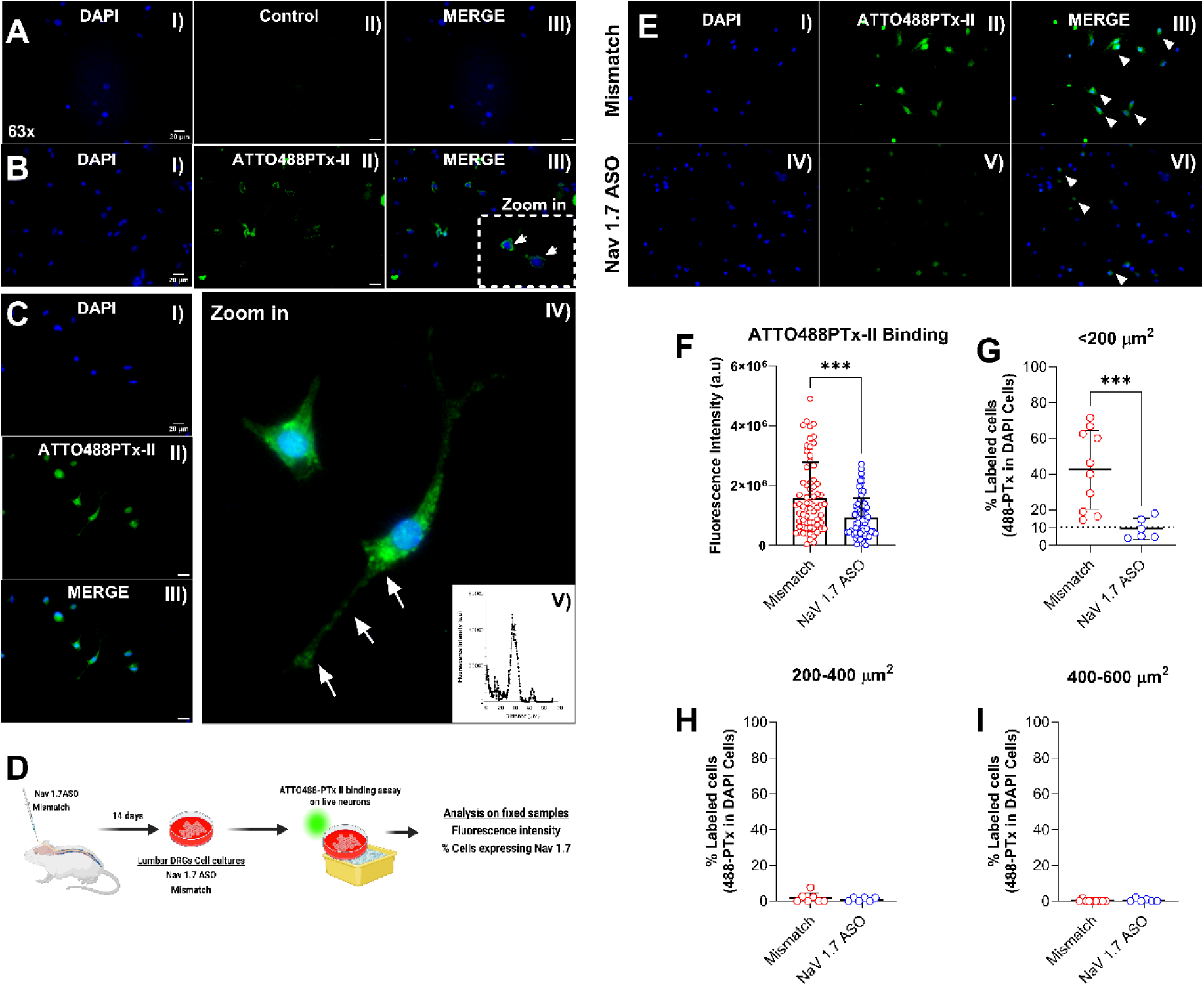
NaV 1.7 antisense reduces the binding of ATTO488PTx-II predominantly in small dorsal root ganglia neurons. Naïve dorsal root ganglia neurons labeled with ATTO488PTx-II display visible fluorescence signal in comparison with non-incubated controls (A-B; I-III). In these naive DRG cell cultures labeled with ATTO488PTx-II (C, I-III) some neurons display punctate signal towards the neuronal soma and axons (C, IV-V, white arrows). The intrathecal injection of NaV 1.7 ASO optimal dose (1000 μg; 10 μL) lead to a significant decrease in binding from ATTO488PTx-II in DRG neurons (F). In small size neurons, the fraction of cells expressing NaV 1.7 channels is significantly reduced in the presence of ASO (G-I). On (F-Mismatch=73 cells; Nav 1.7 ASO=72 cells); On (G-I, Mismatch group n=11 fields analyzed; DAPI cells = 236; 488 PTx II cells = 65; Nav 1.7 ASO group, n=6 fields analyzed; DAPI cells = 786; 488-PTx II cells = 66). Results shown as mean ± standard deviation. Symbols *** indicate P<0.001 for comparison between groups. On (F) Mann Whitney test. On (G) T-student test.

### 3.10. Reduced formalin flinching induced by Nav 1.7 antisense covaries with decrease in ATTO488PTx-II binding at dorsal root ganglia but not at sciatic nerves

The injection of formalin in mismatch animals led to a biphasic flinching response (Figure 6A-B). This nociceptive behavior was significantly affected by the ASO optimal dose at second phase (1000 μg/10μL; Figure 6A-B). The lower ASO dose (300 μg/10μL) had similar profile as mismatch controls (Figure 6A-B). In the whole cell z-stacks analyzed, the mismatch group displayed significant increase in NaV 1.7 binding at ipsilateral DRGs (Figure 6C-IV) in comparison with contralateral DRG controls (Figure 6C-I, and D). This binding was significantly decreased in ipsilateral DRGs by ASO optimal dose (Figure 6CIII/VI, D) but not by the lower dose tested (Figure 6CII/V, D). Interestingly, although different outcomes were observed for NaV 1.7 binding, both doses of ASOs provided similar knockdown of *Scn9a* mRNA at lumbar DRGs and cord (Figure 6E, F). Of note, changes in motor function for these 2 doses were observed, however only during a 24-hour period after ASO delivery, and absent from day 1-14^th^ after injections (supplementary figure 4A-B).

**Figure 6.**
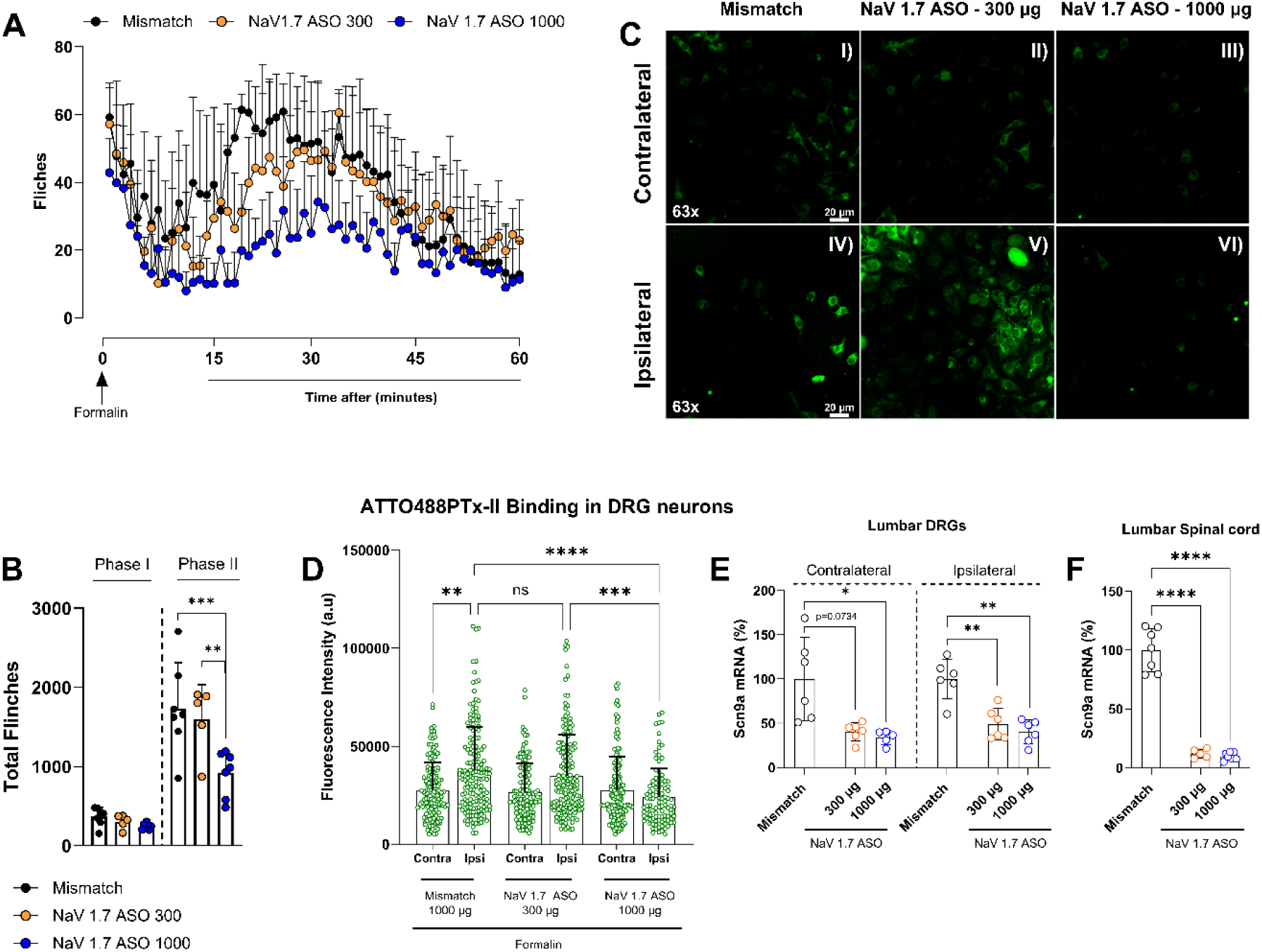
NaV 1.7 antisense affects formalin behavior in a dose-dependent way with different profile for ATTO488PTx-II binding. The lumbar intrathecal delivery of NaV 1.7 antisense (300 or 1000 μg; 10 μL) using spinal catheters led to a different response in formalin nociceptive behavior. The injection of formalin (2.5%; 50 μL; intra-dermal) at the hind-paw led to flinching response (A) with an increase in ATTO488PTx-II binding in ipsilateral DRGs from Mismatch animals. A similar profile is followed by the NaV 1.7 antisense smaller dose (A-D). The optimal dose of NaV 1.7 ASO (1000 μg; 10 μL) was able to reduce the formalin induced flinching at phase II (A, B) and decrease ATTO488PTx-II binding in ipsilateral DRGs in comparison with Mismatch and the smaller antisense dose (C). At lumbar DRGs (contralateral and ipsilateral) and at the lumbar spinal cord (E, F) a significant reduction in *Scn9a* mRNA is seen for both NaV 1.7 antisense doses in comparison with mismatch controls. Results shown as mean ± standard deviation. On (D) Mismatch (Contra = 147 cells/Ipsi = 161 cells), NaV 1.7 ASO 300 μg (Contra= 125 cells/Ipsi=166 cells), Nav 1.7 ASO 1000 μg (Contra = 147 cells/Ipsi = 101 cells). Symbols *, **, ***, and **** indicate P<0.05, P<0.01, P<0.001 and P<0.0001 for comparison between groups. On (B) Two-way ANOVA with Tukey post-hoc test. On (D) Kruskal-Wallis test with Dunn’s post-hoc test. On (E, F) One-way ANOVA with Dunnet’s post-hoc test.

As opposed to what was noticed at the DRG level, the NaV 1.7 antisense treatment was not able to decrease the 488PTx-II binding in the sciatic nerves (Figure 7A-E). Both ASO doses tested do not interfere with binding of PGP 9.5-control (Figure 7AI-III, and C) and 488PTx-II (Figure 7 BI-III, and D) in comparison with mismatch controls. In addition, no changes were detected in *Scn9a* mRNA (Figure 7E).

**Figure 7.**
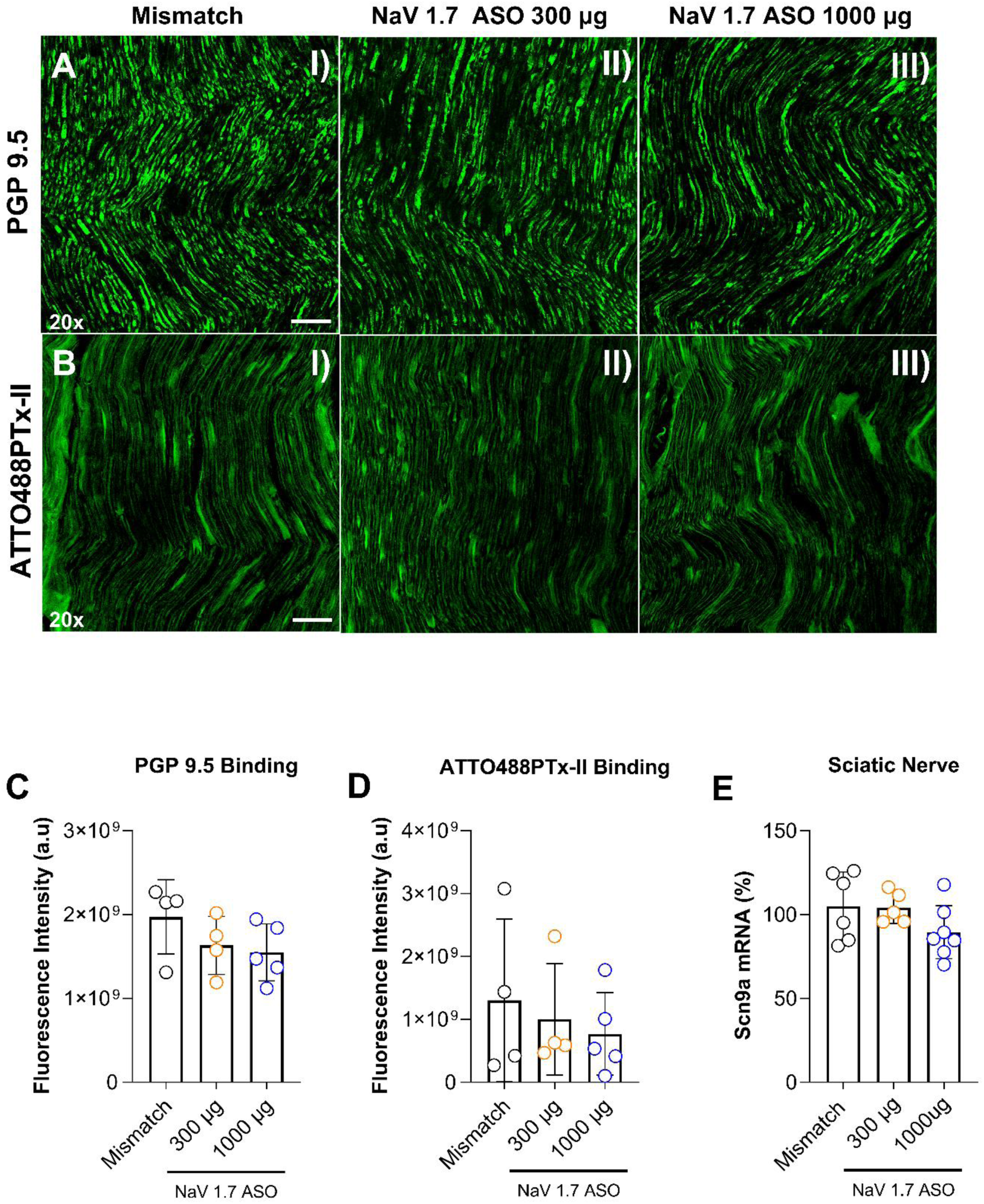
NaV 1.7 antisense does not decrease ATTO488PTx-II binding in sciatic nerves. The lumbar intrathecal delivery of NaV 1.7 antisense (300 or 1000 μg; 10 μL) using spinal catheters did not lead to a decrease in binding of a ATTO488PTx-II binding in sciatic nerves harvested from rats 24 hours after being ipsilaterally stimulated with formalin (2.5%;50 μL; intra-dermal) in the hindpaw (A, B, I-III; D). The *Scn9a* mRNA at the sciatic nerve (E) was not affected by the antisense treatment with both doses tested. Results shown as mean ± standard deviation. On (C, D) Kruskal-Wallis test with Dunn’s post-hoc test. On (E) One-way ANOVA with Dunnet’s post-hoc test.

## 4. DISCUSSION

The relevance of NaV 1.7 for nociceptive processing comes from human studies reporting pain sensitivity covarying with gain or loss of function mutations associated with *Scn9a* gene (4, 42–48). These findings have been recapitulated in pre-clinical studies where NaV 1.7 functionality has been linked with inflammatory and neuropathic pain phenotypes (10, 28, 40, 48–53). In the present studies we undertook a systematic assessment of the dose dependent effects of an intrathecal oligonucleotide targeting NaV 1.7 on pain behaviors, mRNA reduction in DRG and spinal cord and effects upon NaV 1.7 binding. The discussion of our findings will be addressed in specific topics below.

### 4.1. ASO Specificity and Effectiveness Profile

The specificity of our intrathecal NaV 1.7 ASO targeting was reestablished *in vitro* and *in vivo* studies where the active ASO but not its mismatch displayed prominent selectivity with specific block of NaV 1.7 message and with no effects at the highest concentrations on the mRNA of other DRG NaV channels (1.1/1.2/1.6 and 1.8/1.9).

Consistent with the previous use of NaV 1.7 loss of function mutations, neuraxial transfection or antisense (54–59), intrathecal NaV 1.7 targeted ASO in the rat, dose-dependently reduced facilitated states of nociceptive processing as assessed by its effects upon phase 2 formalin flinching, mechanical thresholds assessed in normal and intraplantar carrageenan-treated animals. Similar doses of intrathecal ASO mismatch were without effect. Importantly, the IT ASO-mediated KO of *Scn9a* mRNA had little or no effect upon acute nociception (e.g. phase 1 of the formalin) or low threshold mechanical motor function (e.g. absence of effect upon light touch, motor function or proprioception). The analgesic effect upon mechanical compression evoked withdrawal thresholds is consistent with the likelihood of a facilitated state arising from a degree of local injury resulting from repeated paw compression. While the block of the acute algogenic effect of intraplantar OD-1 in the IT ASO-treated-rat is consistent with a KO of the target binding site, we did not see a loss of NaV1.7 binding in the peripheral nerve. An alternative explanation is that OD1 activated the afferent but the loss of NaV1.7 in the central terminal (or dorsal horn) prevented the nociceptive afferent traffic to the second order neuron (60). The duration of these IT ASO effects is typically long-lasting and were not systematically examined by changes in threshold out to at least 4 weeks were observed.

In our hands, male and female rats treated with the NaV 1.7 ASO optimal dose (1000 μg/10 μL) both exhibited substantial increases in mechanical thresholds assessed through Randall-Selitto apparatus. These observations covary with our ASO uptake findings at the DRG, where ASO particles were detected in large and small neurons. Both cells are known to express NaV 1.7 (5) and contribute to mechanical pain associated with deep-pressure stimulus (10, 40).

The delivery in rats by IT catheters of the optimal NaV 1.7 ASO dose having significant effects upon *Scn9a* mRNA and producing analgesia was well tolerated. Of note, following the ASO, but not the mismatch, while there was unexpectedly a transient dose-dependent reduction in rotarod performance, no effects were observed on the hind paw acute thermal response and there were no changes in body weight over the following 14 days (as shown in supplementary figure 12A-D). In these rats there were few or no changes in IBA-1 (microglia) or GFAP (satellite cells/astrocytes) mRNA in comparison with mismatch and PBS controls (as shown in figure 1F, and G). Notably, previous reports have reported the moderate appearance of DRG and spinal pathology secondary to the catheter placement (61, 62).

### 4.2. Neuraxial ASO distribution and uptake

Neuraxial CSF represents a route for intrathecal solutes to redistribute. However, this space, while subject to local pressure gradients, presents as a poorly distributed volume with bulk rostro-caudal reflective of injection volumes (63). Assessment of the presence of ASO particle revealed that intrathecal delivery of both mismatch or NaV 1.7 targeted antisense lead to rostral distribution of ASO particles after bolus delivery showing a progressive gradient with lower levels at the thoracic and cervical levels relative to the lumbar cord and DRG, and with evident uptake in cells in brainstem and forebrain structures including the cerebellum, cerebral cortex, hippocampus, thalamus (TH) and olfactory bulbs. The intrathecal ASO particles in the intrathecal fluid displayed movement along the root sleeves with cellular uptake into large and small DRG neurons(64). At the spinal level the pia is present as an exclusion barrier for larger molecules and solutes (e.g. high molecular weight dextrans and adenoviruses) (65). As indicated here, given their presence in the spinal parenchyma, the pia does not present such a barrier for these low molecular weight ASOs (66, 67). In tissue, ASOs associate with specific cell surface membrane proteins which mediate internalization, and such productive internalization is observed in both glial and neuronal cell types(68, 69).

### 4.3. Localization of intrathecal ASO effects upon NaV 1.7 expression

The rostro-caudal gradient of ASO distribution resulting from the lumbar bolus delivery covaried with the observed reduction in the *Scn9a* mRNA observed in DRG for any given dose. Thus, as indicated in Figure 1, the greatest silencing occurred at the thoracic lumbar level of injection with progressive reduction in the target DRG effects showing a progressive reduction at rostral levels with the smallest reduction in *Scn9a* mRNA noted at the trigeminal ganglia. Importantly, increasing the intrathecal ASO dose (with a fixed volume resulted in a greater rostro-caudal distribution of NaV 1.7 reduction. These results recapitulate distributional transfection studies after IT AAV injection in the rat model with a lumbar catheter and in the mouse after a percutaneous lumbar bolus, where a robust lumbar to cervical gradient of DRG transfection was observed (54, 70). Of note, in these models with a fixed injection volume the rostro-caudal degree of transfection was increased as a function of concentrations (e.g. viral titer). An important issue that is not often considered is that the *Scn9a* message is observed in the dorsal horn. While it has been reported that functional NaV 1.7 are present in substantia gelatinosa neurons, the source of these channels is speculative, being suggested that there is transport from the afferent terminal to the second order neurons (6). However, others have also reported low levels of NaV 1.7 message in spinal dorsal horn (71, 72). We note here that the dorsal horn message was effectively abolished by even low levels of IT NaV 1.7 ASO, which suggests potential effects upon non-afferent neuraxial components occuring as a result of movement of the ASO though the pial barrier.

### 4.4. ASO dose effects on pain behavior

In the present studies, we show that the intrathecal injection of antisense oligonucleotides in doses between 100 and 3000µg, led to a dose-dependent silencing of *Scn9a* mRNA in lumbar (L1-L5) dorsal root ganglia harvested at 14 or 28 days after the single bolus ASO delivery at spinal level in rats. In all behavioral models, the lowest IT ASO dose that resulted in a significant reduction in the measured nociceptive behavior as assessed by high intensity mechanical compression, phase 2 formalin flinching, and inflammation induced tactile allodynia, was 1000µg. Assessment of the relationship between *Scn9a* message reduction and intrathecal ASO dose similarly revealed a close covariance with the maximum reduction in the Lumbar DRG being in the range of 50 - 75%. We note that for doses such as 100 and 300, the degree of message suppression was often similar, but the analgesic effects were limited to the higher dose concentrations. This observation brings to the fore the appreciation that increasing the injected dose by increasing concentration (vs increasing injection volume) not only increases the local tissue exposure at the injection site but also results in delivering a higher solute concentration to targets that lie more distal to the injection level, that may contribute to the nociceptive signal underlying the pain state. It is important to note that for models such as formalin, evidence of terminal activation as measured by, for example, C-fos leads to greatest activation in the L3-5 segments but significant expression may be observed at segments ranging from S1-T13. Further, in acute models the primary activation is superficial (e.g the substantia gelatinosa), whereas in more chronic exposures the activation shifts to the deep dorsal horn (73–75). So, the concentration at the local DRG site may be maximum with a given dose (concentration) but when volume of delivery remains fixed, the higher concentrations may lead to a broader rostro-caudal distribution of NaV 1.7 knock-out and further affect nociceptive signaling. This organization may reflect upon the role played by NaV 1.7 expressing circuits in the spinal parenchyma.

### 4.5. Quantitative assessment of NaV 1.7 antagonist binding

ATTO488PTx-II (488-PTx II) is a NaV 1.7 antagonist that carries the ATTO488 azide tag. This allows fluorescence microscopy imaging at 488nm (19). This toxin is highly selective for NaV 1.7 binding at the external voltage sensing site, displaying minimal affinity for other NaV Channels (19). To further address this issue, we examined (as shown in Figure 4) the covariance of signals in HEK cells transfected with NaV 1.7-Halotag channels and 488-PTx II. In the ex*-vivo* studies, consistent with preferential binding to nociceptors, the 488-PTx II binding in DRG cultures showed reliable up take in small (<200 µm^2^) but not larger cells which are not typical nociceptors in the rodent (76). In our binding studies with the rat DRG cell culture system, we observed that knockdown for *Scn9a* gene, using the optimal intrathecal ASO dose, displays a significant reduction in 488-PTx II binding in comparison with mismatch controls, both as measured by fluorescence intensity (reflecting binding in cells still expressing the channels) and the number of cells expressing NaV 1.7 binding, suggesting that some cells experience a full knockout of the channel. These data confirm the utility of the 488-PTxII binding in defining the presence of membrane expression NaV 1.7 channels, while the ASO studies with knockdown confirm the importance of NaV 1.7 in its functionality.

### 4.6. Covariance of analgesia with ASO doses, mRNA and Binding

We sought to systematically assess the binding/message covariance using the intrathecal delivery of the NaV 1.7 ASO in the hind paw formalin flinching model. Here we observed that the IT 1000µg -group displayed a highly significant suppression of phase 2 flinch as compared to the mismatch or the 300µg-NaV 1.7 ASO. In parallel, the ipsilateral NaV 1.7 binding on DRGs of the 1000 µg dose, but not the 300 µg rats, was significantly reduced as compared to mismatch controls. Strikingly, in these groups the ipsilateral reduction in NaV 1.7 message were both significantly reduced in comparison to the mismatch. This difference between message and channel expression suggests that the binding expression presents as a closer covariance to the actual analgesic variable than the message expression itself. One explanation may reflect the time required for turnover of the membrane express channels (as we believe to be measured by the 488-PTx II binding). As noted, these studies examined binding and mRNA at 14 days and studies on NaV turnover suggest that this event may reflect intervals of several weeks (77). How the ASO dose would impact NaV membrane turnover is not clear. It is notable that the NaV 1.7 channel displays turnover, and its replacement involves the role of a transporter protein (CRMP2) (78), although how these ASOs reflect such transport is not known. A second possibility may be that the lower ASO dose, as discussed above, may have resulted in a more limited rostro-caudal distribution of target engagement and that the lack of effect at the lower ASO dose reflects a lack of KO at distal DRGs or parenchymal tissue that is contributing to the formalin flinching.

### 4.7. Effects of the pain state on knock down

We observed a more effective knock-down of *Scn9a* mRNA by the ASO optimal dose (1000 μg/10μL) in DRGs at inflammatory pain triggered by carrageenan insult rather than formalin. These different profiles of knock-down varying according to stimulus may reflect different degrees to which NaV 1.7 signaling is regulated. In addition, although we performed the analysis from DRG-ASO uptake only 14 days after intrathecal delivery, the differences in knock-down profile seen in DRGs harvested in different endpoints also reflect a kinetic relationship between the bioavailability of oligonucleotides in the cell compartment and regulation of target mRNA. Assuming a drug concentration paradigm, such differences seen for antisense knock-down between day 14 and day 28 endpoints suggest that the ASO would better regulate transcriptional alterations involving *Scn9a* at 14 days rather than 28 days, and thus effectively mitigate pain behavior depending on NaV 1.7 early on.

### 4.8. Skin and ASO effects

Although heavily expressed in neuronal cells, a series of voltage gate-sodium channels can also be found in rats and human keratinocytes, such as NaV 1.7 (79). In formalin test, data suggests that keratinocytes can be activated by formalin and release calcium stores from endoplasmic reticulum (80), which further triggers the release of several molecules that can alter neuronal excitability, such as ATP, Prostaglandin E_2_, and NGF (81, 82). Indeed, the ratio of drug in the bloodstream after intrathecal administration of ASO is small if compared with the parenteral or subcutaneous routes typically used for antisense drug administration (83). Still, a possibility that needs to be considered is given the higher concentration provided by the 3000 μg ASO dose, a small portion of antisense particles could reach systemic distribution and target other cell types carrying *Scn9a* gene such as keratinocytes in the epithelial tissue (79). The increase in NaV 1.7 binding ipsilaterally at mismatch controls reflects the ability of neuropeptides typically released at peripheral tissue by formalin, such as CGRP, Substance P, and Serotonin to act in peripheral terminals and increase NaV 1.7 signaling at DRG neurons. Of note, the nociceptive activity of these 3 neuropeptides is reported to decrease in NaV 1.7 knockouts (10, 84) or by specific blockers (85).

### 4.9. Peripheral Nerve NaVs and ASO effects

A point of interest was the observation that NaV 1.7 binding was not lost in peripheral nerve 14 days after IT ASO. It has been suggested that axons from primary afferent neurons may display local biosynthesis of some voltage gate sodium channels, such as NaV 1.6 (86). This biosynthesis is performed by the endoplasmic reticulum (ER) - Golgi route, where channels are stored at the ER and subsequently transported to the axonal membrane (86). To determine if the ASOs induced knockdown of *Scn9a* mRNA in additional sites that could be involved with neuronal protein synthesis (e.g., sciatic nerves and terminals), we assessed the effects of the intrathecal ASO dosing of 1000 and 3000μg doses on intraplantar induced OD-1 flinching. Both ASO doses prevented the OD-1 nociceptive response, with no differences between them regarding *Scn9a* mRNA at spinal cord and DRG level. Thus, at least for the 3000 ug ASO dose, the possibility of additional knockdown outside the DRG structure could not be suggested as a mechanism to explain the different pharmacological outcomes provided by this dose. Such studies will be of interest to fill the gaps related to the trafficking of these channels in primary afferent neurons at healthy and pathological phenotypes.

## 5. CONCLUDING REMARKS

The present studies demonstrate the efficacy of IT ASO selectively targeting neuraxial NaV 1.7 in producing a robust dose-dependent analgesia as measured in models of facilitated processing. Our studies assessing membrane expression of the channel using a fluorophore labeled ligand binding at an extracellular site on the NaV 1.7 protein revealed the importance of assessing the expression of the membrane expressed channel vs mRNA. The presence of message in the spinal parenchyma raises the possibility that a component of the efficacy of the NaV 1.7 reduction reflects upon these effects on preventing the role of NaV 1.7 in the dorsal horn systems. In that event, it is noteworthy that unlike other platforms affecting spinal protein expression, ASOs can penetrate the pial barrier to block transcription.

## 6. AUTHOR CONTRIBUTIONS

Kaue Franco Malange: conceptualization, methodology, investigation, data curation, data analysis, writing original draft. Julia Borges Paes Lemes: methodology, investigation, writing original draft. Briana Noble: methodology, investigation, data curation, data analysis, writing original draft. Yuvicza Anchondo: methodology and investigation. Carlos Morado Urbina: methodology, data curation, data analysis, writing original draft. Saee Jadhav: methodology and data curation. Sara Dochnal: formal analysis and writing-review and editing. Chiraag Kambalimath: methodology and data curation. Pedro Alvarez: conceptualization, methodology, investigation, data curation, and data analysis. Bethany Fitzsimmons: conceptualization, methodology, investigation, data curation, data analysis, and writing-review and editing. Curt Mazur: conceptualization, methodology, investigation, data curation, and data analysis. Holly B. Kordasiewicz: conceptualization, methodology, investigation, data curation, and data analysis. Kim Dore: conceptualization, supervision, writing-original draft, writing-review and editing. Hien T. Zhao: supervision, project administration, resources, writing-review and editing. Tony Yaksh: conceptualization, supervision, project administration, resources, writing-original draft, writing review and editing.

## SUPPLEMENTAL FIGURES

**Supplemental figure 1.**
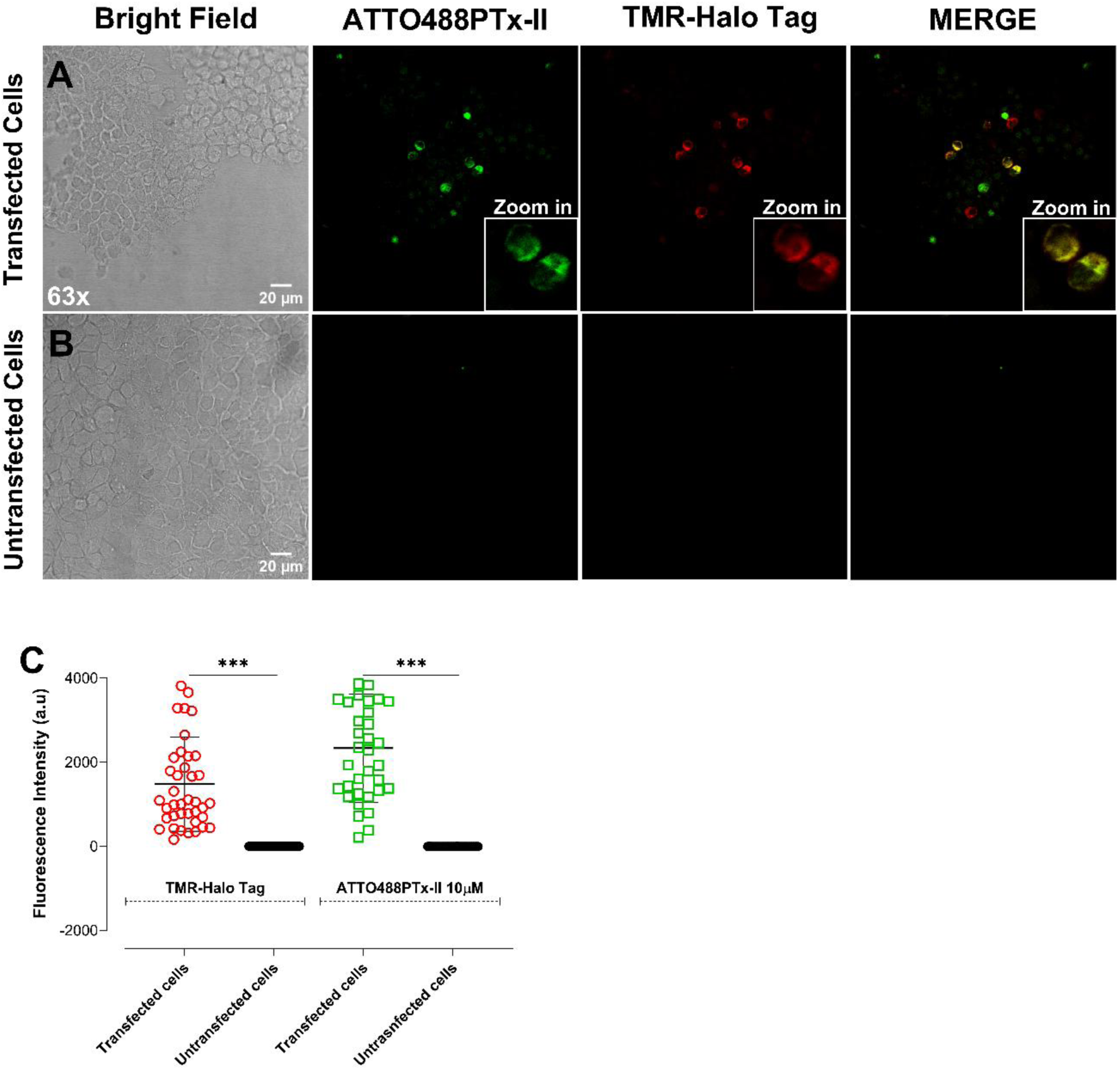
HEK cells expressing NaV 1.7 channels significantly showed fluorescent signal in the presence of TMR-Ligand at 2.5 μM, and 488-PTx II at 10 μM dose. **(A)** HEK cells transfected with NaV 1.7 channel exhibited fluorescence signal in the presence of 488-PTx II and TMR-Ligand, indicating a selective binding from the channel antagonist which was confirmed by the colocalization at the Merge field. **(B)** No signal was detected in un-transfected cells for 488-PTx II and TMR-Ligand suggesting minimal or none off-target molecular effects. **(C)** The fluorescence intensity of transfected cells exposed to 488-PTx II and TMR-Ligand is significantly different in comparison with un-transfected cells. Results shown as mean ± standard deviation. Symbol *** indicates P<0.001 for comparison between groups. Welch’s T-test.

**Supplementary figure 2.**
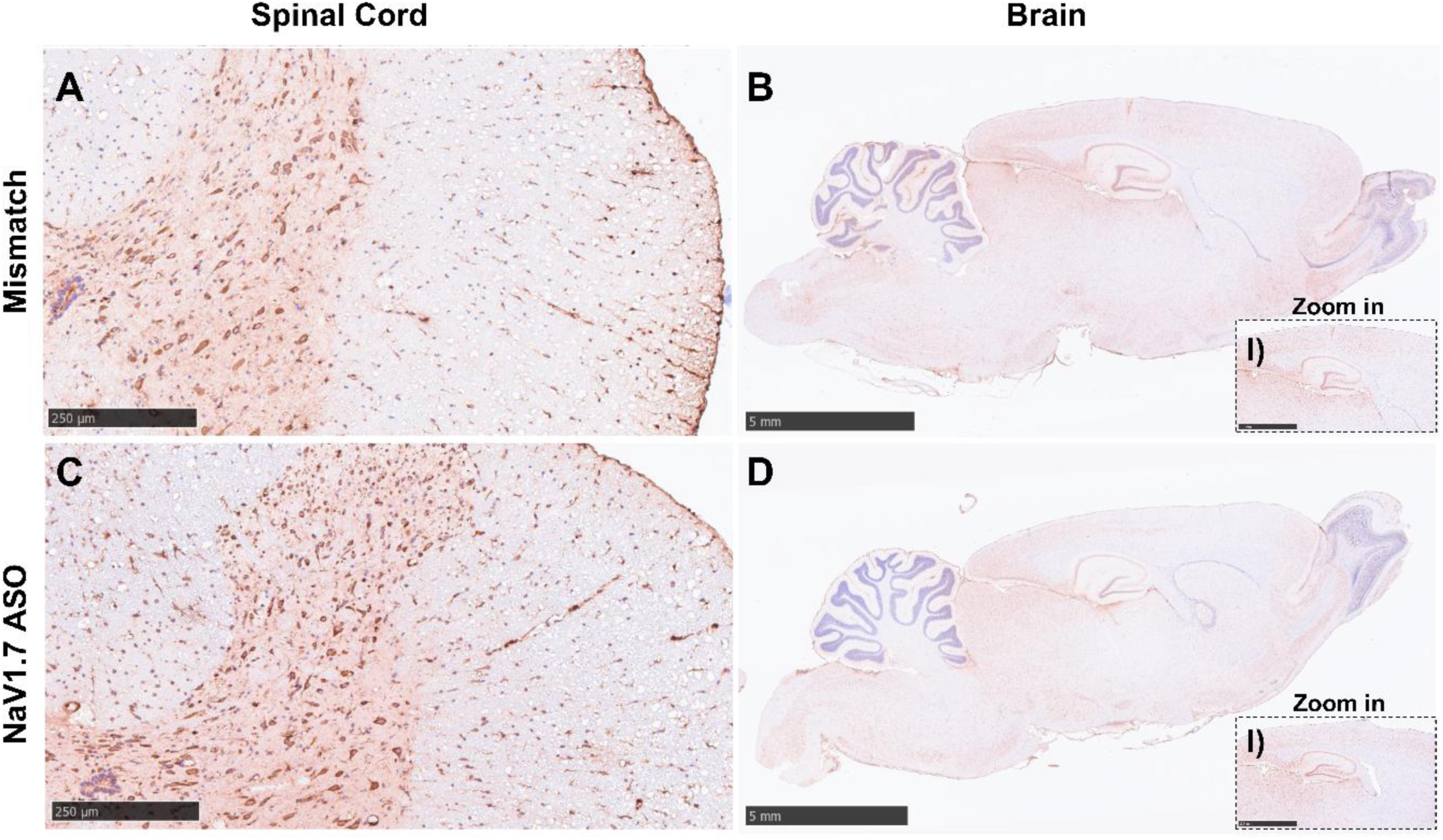
Antisense oligonucleotides are uptake at the spinal cord and brain. The lumbar intrathecal delivery of Mismatch or NaV 1.7 ASO antisense using spinal catheters lead to rostral distribution and antisense uptake at thoracic section of the spinal cord (A-C) and Brain (B, D; B-I and D-I for higher magnification). The thoracic spinal cords and brain were harvested 14 days after the intrathecal delivery of ASOs. Pictures were taken at magnification of 7.5x, 0.5x and 1x respectively on (A)/(C), (B)/(D), and (B-I)/(D-I).

**Supplementary figure 3.**
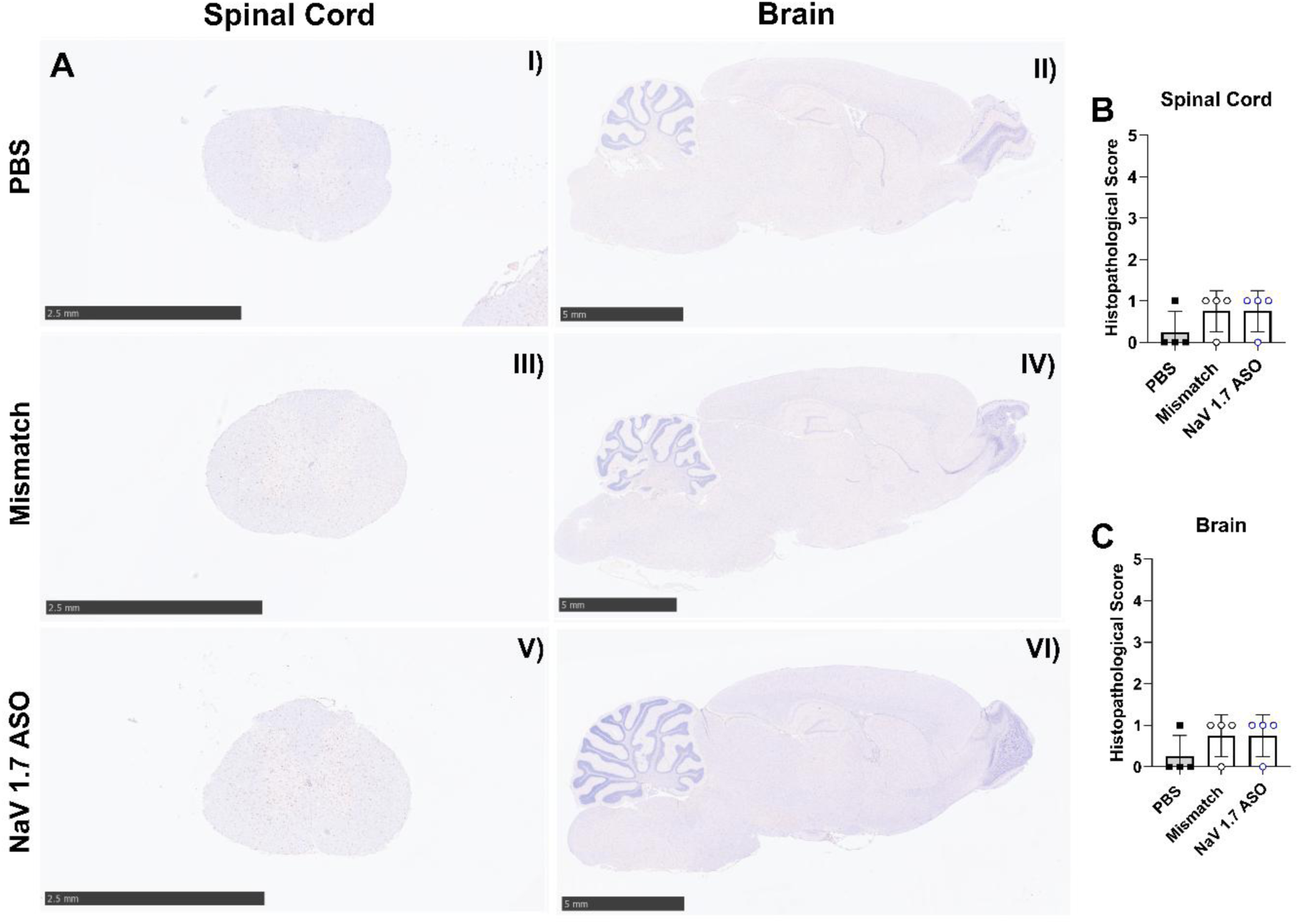
Assessment of spinal cord pathology after intrathecal antisense delivery. The intrathecal injection of Mismatch or NaV 1.7 ASO optimal dose (1000 μg; 10 μL) lead to none to slight microgliosis (IBA-1 staining) at the spinal cord in comparison with PBS controls (A-I, IIII, and V; and B for histopathological score). In similar way, this was also observed at the brain (A-II, IV, and VI; and C for histopathological score). However, no these observational changes did not significant impact the score between the groups analyzed (B/C). Results shown as mean ± standard deviation. Kruskal-Wallis test with Dunn’s post-hoc test.

**Supplementary figure 4.**
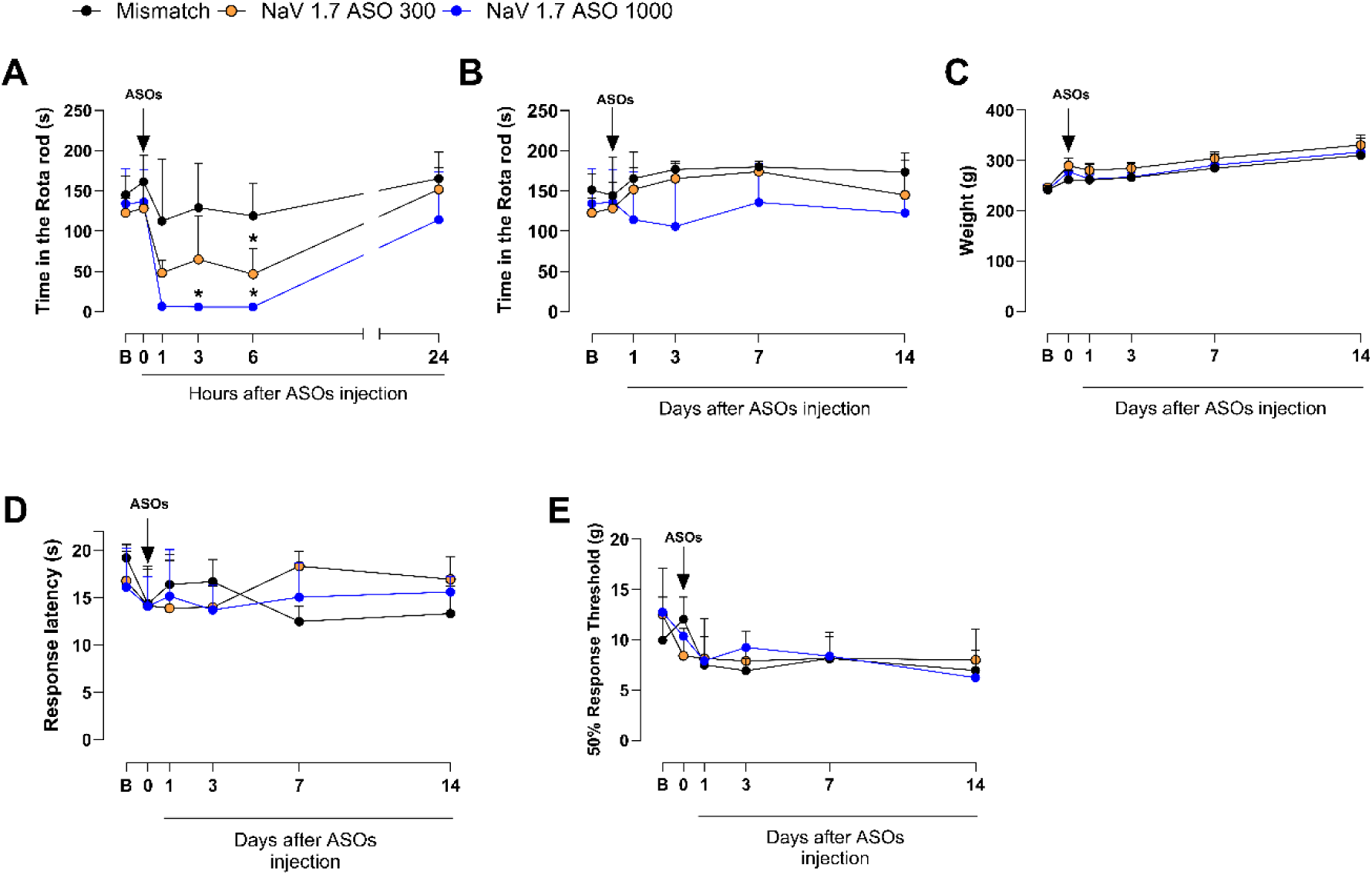
Assessment of motor function, tactile and thermal thresholds from different doses of ASO. The intrathecal injection of ASO (300 μg/10μL or 1000 μg/10μL) led to a significant decrease in rota-rod activity at 3, 6h after ASO delivery **(A)**. This impairment on motor function is not seen day 1-day14 after ASOs injection **(B)**, where animals also show normal weight-gain **(C).** No changes in thermal (D) or tactile (E) thresholds were detected between the 2 doses tested in comparison with mismatch controls. Results shown as mean ± standard deviation. Symbol * indicate P<0.05 in comparison with mismatch controls. From A-E, Two-way ANOVA with Tukey post-hoc test.

